# Sex and BDNF Val66Met Polymorphism matter for exercise-induced increase in neurogenesis and cognition in middle-aged mice

**DOI:** 10.1101/2022.07.29.502070

**Authors:** Dannia Islas-Preciado, Tallinn F.L. Splinter, Muna Ibrahim, Natasha Black, Sarah Wong, Stephanie E Lieblich, Teresa Liu-Ambrose, Cindy K Barha, Liisa A.M. Galea

## Abstract

Females show greater benefits of exercise on cognition in both humans and rodents, which may be related to brain-derived neurotrophic factor (BDNF). A single nucleotide polymorphism (SNP), the Val66Met polymorphism, within the human *BDNF* gene, causes impaired activity-dependent secretion of neuronal BDNF and impairments to some forms of memory. We evaluated whether sex and BDNF genotype (Val66Met polymorphism (Met/Met) versus wildtype (Val/Val)) influenced the ability of voluntary running to increase cognition and hippocampal neurogenesis in mice. Middle-aged C57BL/6J (13 months) mice were randomly assigned to either a control or an aerobic training (AT) group (running disk access). Mice were trained on the visual discrimination and reversal paradigm in a touchscreen-based technology to evaluate cognitive flexibility. BDNF Met/Met mice had fewer correct responses compared to BDNF Val/Val mice on both cognitive tasks. Female BDNF Val/Val mice showed greater cognitive flexibility compared to male mice regardless of AT. Despite running less than BDNF Val/Val mice, AT improved performance in both cognitive tasks in BDNF Met/Met mice. AT increased neurogenesis in the ventral hippocampus of BDNF Val/Val mice of both sexes and increased the proportion of mature type 3 doublecortin-expressing cells in the dorsal hippocampus of female mice only. Our results indicate AT improved cognitive performance in BDNF Met/Met mice and increased hippocampal neurogenesis in BDNF Val/Val mice in middle age. Furthermore, middle-aged female mice may benefit more from AT than males in terms of neuroplasticity, an effect that was influenced by the BDNF Val66Met polymorphism.

**Highlights:**

- BDNF Met/Met mice performed worse than BDNF Val/Val mice in middle-age
- Aerobic training (AT) increased cognitive performance in BDNF Met/Met mice
- AT increased neurogenesis in middle-aged BDNF Val/Val mice only
- Female BDNF Val/Val mice had better cognitive flexibility than males regardless of AT
- AT increased more mature new neurons in middle-aged female mice

---

Normal aging is associated with reductions in a number of cognitive domains, including episodic memory and executive functions (Levine et al., 2021). Coupled with cognitive decline are changes in the brain, notably the hippocampus, a region that is important for learning and memory (Sang et al., 2021). Hippocampal volume decreases with age, which is associated with higher risk for cognitive impairment and dementia (Raz et al., 2005; Jack et al., 2010). Sex differences in cognitive decline with normal aging are seen as human females have greater cognitive decline in executive functions with age (Levine et al., 2021). It is crucial to find a viable, non-invasive and feasible strategy to prevent decline and improve cognition in older age. Recently the World Health Organisation suggested engaging in physical activity to slow down cognitive decline (*Risk Reduction of Cognitive Decline and Dementia*, 2019). However, given the sex differences seen in cognitive decline with normal aging and in dementia risk (Irvine et al., 2012; Levine et al., 2021), it is important to understand how sex may affect the role of exercise to offset cognitive decline with aging, particularly in middle-age as an important time for intervention (Barha and Liu-Ambrose, 2020).

Physical exercise, in the form of aerobic training (AT), promotes cognition including prefrontal cortex (PFC)-dependent executive functions, and increases hippocampal volume in older adults with or without mild cognitive impairment (MCI; a prodromal state to Alzheimer’s disease(AD)) (Baker et al., 2010; Liu-Ambrose et al., 2018; Erickson et al., 2011; Makizako et al., 2015; ten Brinke et al., 2015). AT also increases hippocampal-dependent learning and memory in rodents, but fewer studies have investigated AT effects on executive functions in rodents (Barha et al., 2017b). Evidence suggests AT effects on cognition may vary by sex, where executive functions in older human females may benefit more than human males from AT (Barha et al., 2017a, 2017b; Middleton et al., 2008). Interestingly, in rodents, a meta-analysis suggested sex differences in cognitive benefits from AT depending on the type of cognition as females showed greater benefits on spatial tasks whereas males showed greater benefits on non-spatial and conditioned avoidance tasks after AT (Barha et al., 2017b). These sex differences may be related to the ability of AT to influence neurotrophic factors such as brain-derived neurotrophic factor (BDNF) (Barha et al., 2019a).

Exercise, sex, and estrogens influence BDNF (Barha et al., 2017a, 2017b; Chan and Ye, 2017). BDNF levels are increased after exercise in humans (plasma and serum) and rodents (hippocampus, corpus striatum and amygdala) (Bastioli et al., 2022; Barha et al., 2017a, 2017c; Erickson et al., 2011; Liu et al., 2009; Marais et al., 2009; Pietrelli et al., 2018; Short et al., 2022; Triviño-Paredes et al., 2016; Voss et al., 2013). BDNF is a neurotrophin strongly involved in the neurogenic process, and modulates the effects of exercise on brain outcomes (Berchtold et al., 2001; Cowansage et al., 2010; Triviño-Paredes et al., 2016; Voss et al., 2013). BDNF levels are also negatively associated with more cognitive impairment in AD (Erickson et al., 2010; Siuda et al., 2017), and positively associated with increased hippocampal volume and spatial memory performance (Erickson et al., 2010; Voss et al., 2013). Sex differences in BDNF levels after exercise have also been noted, with human and rodent females showing a greater increase in BDNF levels after exercise compared to males, which are related to ovarian hormone levels (Barha et al., 2017a, 2017b, 2017c; Berchtold et al., 2001).

A single nucleotide polymorphism (SNP) is found within the pro-domain region of the human BDNF gene, resulting in an amino acid substitution of valine (Val) to methionine (Met) at codon 66, termed the Val66Met polymorphism. The BDNF Val66met polymorphism reduces activity dependent neuronal secretion of BDNF (Egan et al., 2003; Miranda et al., 2019; Park et al., 2017), and a robust body of literature suggests it is associated with impaired hippocampal-dependent memory (Chen et al., 2008; Dincheva et al., 2014; Egan et al., 2003; Kennedy et al., 2015; Lamb et al., 2015). However, less is known about other cognitive domains such as PFC-dependent executive functions and, intriguingly, BDNF Met/Met have compromised PFC activity (Pattwell et al., 2012). Under the umbrella term for PFC-dependent cognition, cognitive flexibility is a part of executive functioning and is a fundamental domain needed in order to adapt our behavior to cope with novelty, and thus facilitate adaptation (Diamond, 2013; Logue and Gould, 2014; Armbruster et al., 2012; Dajani and Uddin, 2015). Notably, cognitive flexibility is one of the first domains affected in pathological and non-pathological aging (Corbo and Casagrande, 2022; Guarino et al., 2020), and impairments in cognitive flexibility could be a predictor of later onset of AD, perhaps due to the early degeneration of the PFC in AD (Traykov et al., 2007; Salat et al., 2001).

The hippocampus is a highly plastic structure including the presence of adult neurogenesis in the dentate gyrus (Christie and Cameron, 2006). Hippocampal neurogenesis exists in all mammalian species, including humans (Christie and Cameron, 2006; Eckenhoff and Rakic, 1988; Eriksson et al., 1998). Although a couple of studies suggest that hippocampal neurogenesis is non-existent or rare in humans (Franjic et al., 2022; Sorrells et al., 2018), the vast majority of studies, using a variety of techniques, show that adult hippocampal does occur in humans (Boldrini et al., 2018; Moreno-Jiménez et al., 2021; Spalding et al., 2013; Zhou et al., 2022). Intriguingly, adult hippocampal neurogenesis decreases with age, is impaired in AD, and increases with exercise. In rodents, exercise increases the expression of doublecortin (DCX), a protein expressed in immature neurons (Brown et al., 2003) in both young (Fuss et al., 2010; Segal-Gavish et al., 2019) and aged male mice (Connolly et al., 2022), and young female mice (Kim et al., 2017; Segal-Gavish et al., 2019), suggesting that exercise increases new neurons. Moreover, hippocampal neurogenesis promotes cognitive flexibility, and is modulated by both sex and BDNF levels (Anacker and Hen, 2017; Berdugo-Vega et al., 2021; Choi et al., 2018; Kuhn et al., 1996; Marrocco et al., 2017; Mu and Gage, 2011; van Praag et al., 1999; Yagi and Galea, 2019). BDNF is required for the survival of new neurons, and increasing BDNF in AD mouse models increases hippocampal neurogenesis and cognition to levels that mimic the positive cognitive effects of exercise on AD (Sairanen et al., 2005; Choi et al., 2018). Sex differences are seen in both hippocampal structure and function, including neurogenesis, (Yagi and Galea, 2019) and exercise may increase neurogenesis to a greater degree in females than in males (Ma et al., 2012; Ransome and Hannan, 2013). Due to the inconsistent findings across studies examining the influence of the BDNF polymorphism on neuroplasticity and cognition, researchers have suggested the importance of age and sex as possible factors that could shed light on the inconsistencies (Tsai, 2018). Thus it is important to consider sex in studies examining the influence of BDNF polymorphisms, exercise and neurogenesis.

In the present study, we examined sex differences in cognitive flexibility and neurogenesis in response to AT in middle-aged BDNF Val/Val and BDNF Met/Met mice. To our knowledge, no previous study has addressed the effect of AT on cognitive flexibility assessed through a translational tool in middle-aged male and female rodents carrying the BDNF val66met polymorphism. Thus, we evaluated the effect of voluntary running in middle-aged male and female BDNF Val/Val and Met/Met mice on cognitive flexibility. We expected that the BDNF Val/Val mice would show superior cognition and neurogenesis, particularly in females, compared to the Met/Met mice and that AT would have differential effects on cognition and neurogenesis based on genotype and sex.

## Methods

### Subjects

C57BL/6J knock-in mice in which the human BDNF Val66Met SNP was inserted into the mouse BDNF gene were obtained via the University of Calgary from a stock that was originally from Francis Lee. Mice were bred and aged to 13 months old in our breeding facilities during which time they were group housed (2 per cage) in a temperature and humidity controlled colony room (21 ± 1 °C; 50 ± 10% humidity), with a 12-h light/dark cycle (lights were on at 07:00 h). A total of n=34 BDNF Val/Val mice (18 males, 16 females) and n=28 BDNF Met/Met mice (13 males, 15 females) were used. Mice were given food (Purina chow) and water ad libitum before assignment to the experimental groups. All experimental procedures were approved by the Animal Care Committee at the University of British Columbia and were performed according to the Canadian Council on Animal Care guidelines.

### Exercise Intervention

Middle-age male and female mice were single-housed in Digital Ventilated Cages^®^ (Techniplast, Italy) with 12-h inverted light/dark cycle (lights were off at 07:00 h) and randomly assigned to the control sedentary or the Aerobic Training (AT) group. Eight groups were formed for sex by genotype by AT intervention as follows: Male BDNF Val/Val control (n=8), male BDNF Val/Val AT (n=10), female BDNF Val/Val control (n=8), female BDNF Val/Val AT (n=8); male BDNF Met/Met control (n=6), male BDNF Met/Met AT (n=7), female BDNF Met/Met control (n=8), female BDNF Met/Met AT (n=7). The AT group was housed with a running wheel (Bio-Serv Igloo fast-trac, circumference 479 mm) and the distance ran was recorded daily. Access to the running wheel was maintained throughout the experiment. Control sedentary mice were housed without a running wheel.

### Behavioral assessment

Mice were food-restricted to 85-90% of their free-fed body weight to prime seeking behavior for food reward to complete the task (based on Buscher et al., 2017). Mice reached target weight in ~7 days and during habituation to the food-restriction regimen, mice were exposed to 8 μl of the strawberry milkshake reward (Neilson Dairy, Saputo Inc.) placed in a small container inside the home cages. To assess cognitive flexibility, mice were trained on the Pairwise (Visual) Discrimination and reversal paradigm of the Bussey-Saksida touchscreen system (Lafayette Instrument, U.K.).

### Apparatus

Touchscreen-based technology is a highly translatable tool to explore cognition among rodents, humans and other species (Izquierdo et al., 2017; Turner et al., 2017). The Bussey-Saksida touchscreen system consisted of four, individual trapezoidal chambers (238mm wide at screen x 170mm tall x 532 mm deep) equipped with a milk receptacle, an infra-red nose-poke detector, a light and tone generator, and an external reward dispenser connected to the reward tray inside the chamber. On the opposite wall to the tray was a touch-sensitive screen covered with a metal template dividing the monitor into two squares to restrict the animal’s choice field. The divided touch-sensitive screen displayed the two stimuli to be discriminated against. The base was composed of non-shock perforated floors with a waste-collector tray underneath. All chambers were placed inside sound-proofing boxes equipped with a small fan and a video camera to record each session. For an Illustration of the touchscreen system see supplemental Figure 3.

### Experimental timeline

The experimental timeline is shown in Figure 1. Mice were randomly assigned to either control sedentary or voluntary AT groups. Mice were allowed two weeks with the running wheels (or no running wheels) before the cognitive training protocol commenced. The running wheel was introduced in the home cage after the two acclimation weeks elapsed. After 3 weeks of voluntary AT, each animal was weighed for 3 consecutive days and then were kept at 85-90% of their free-feeding weight. Milk reward was introduced in their home cages before behavioral training commenced. Mice were trained for approximately 15 days and then promoted to the visual discrimination acquisition phase. Mice learned to visually discriminate the stimulus associated with the milk reward for a total of 13 sessions (1 session/day). Finally, mice were subjected to the reversal paradigm for 14 days. Twenty-four hours after the last behavioral session, mice were perfused and brains were extracted. Blood (after aortic cut via the thoracic cavity) and adrenal glands were also collected (see below).

**Figure 1.**
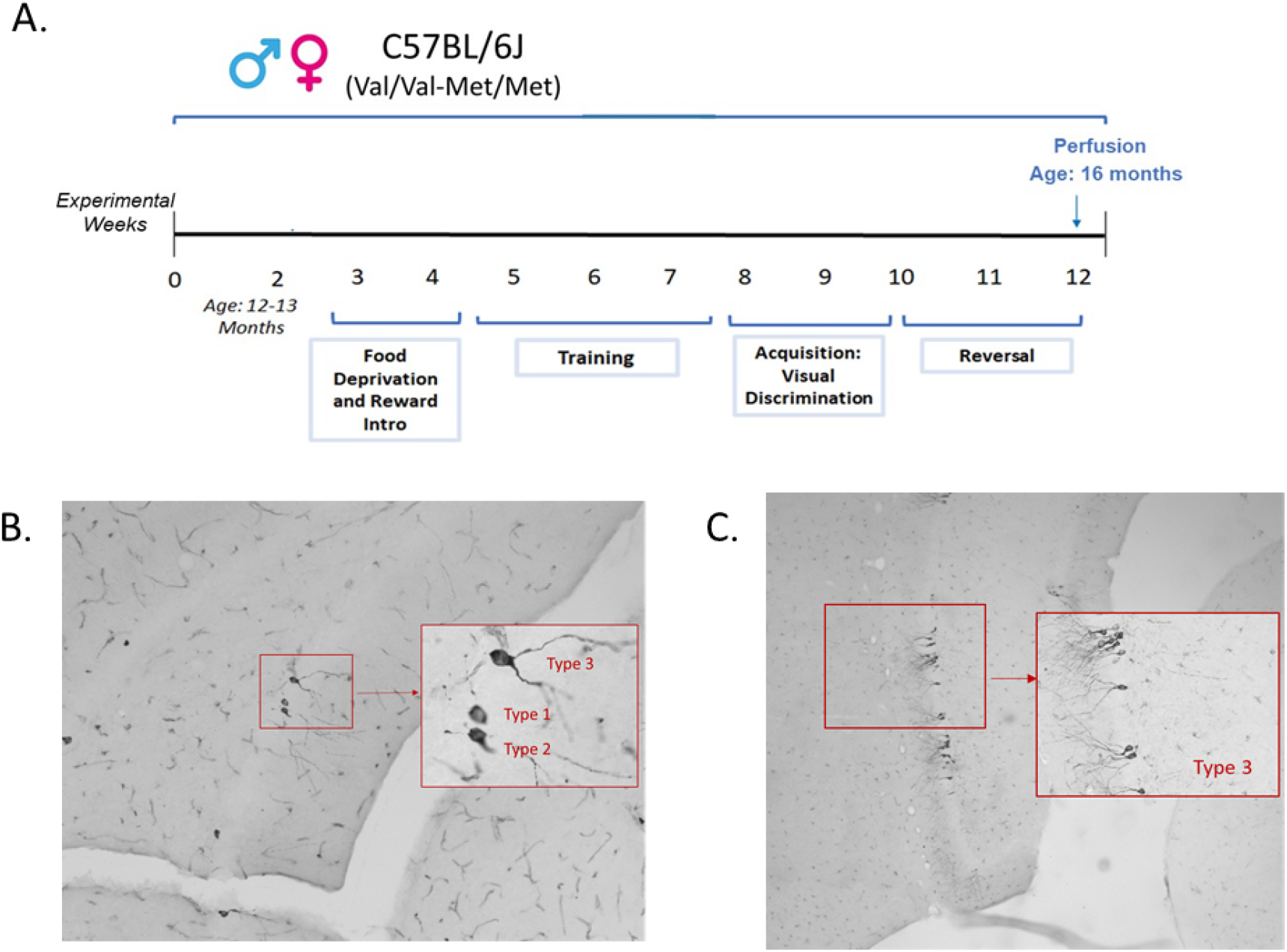
A) Experimental timeline of C57BL/6J mice (with either the brain-derived neurotrophic factor (BDNF) Met/Met genotype or the BDNF Val/Val genotype), behaviour and testing. B) Large photomicrograph (taken with a 20x objective) of an example of a type 1, type 2 and type 3 doublecortin (DCX) expressing cells from a female mouse. Inset was taken with a 40x objective. C) Photomicrograph (taken with a 20x objective) of type 3 DCX expressing cells from a female mouse. Inset was taken with a 40x objective. Representative DCX immunohistochemistry images from each group are shown in supplemental Figure 2.

### Training protocol

#### Pre-training

The pre-training protocol served to train mice how to operate the touchscreens, and was performed with minor changes to Lafayette Instrument’s manual in the following 5 stages: (i) habituation (A and B), (ii) initial touch, (iii) must touch, (iv) must initiate, and (v) punish incorrect. After reaching criterion for a particular stage, each mouse progressed to the next pretraining stage in the following session. To avoid over-training, upon reaching criterion for ‘punish incorrect’, mice were given 2 days rest followed by a reminder session until all subjects reached criterion at 70% [based on Horner et al., 2013]. In this way, all animals began Visual Discrimination training on the same day. Pretraining took 15-22 days.

#### Visual Discrimination Acquisition

After pre-training completion, visual discrimination training began as mice learned to select the correct stimulus displayed on the screen to elicit the reward delivery response. The trial began with the display of two novel stimuli, one on each side of the screen. The task requires mice to discriminate between these stimuli and to learn which stimulus is “correct”, or associated with the reward delivery, (S^+^) and which one was “incorrect” (S^-^) and not associated with reward. Placement of S^+^ on the screen was determined pseudorandomly, as the S^+^ image would not appear on the same side more than three consecutive times. S^+^ and S^-^ images were counterbalanced across groups. The images used were “marble” and “fan” (see Figure 1), as this pair of pictures is recommended for the Visual Discrimination and Reversal task in mice (Horner et al., 2013). The mouse must ‘nose poke’ the S^+^ image to elicit the strawberry milk delivery (dispensed from the tray), which was accompanied by the illumination of the reward-tray light and a tone. Once the tray was entered and the reward collected, the tray light turned off and a 10 sec ITI began, followed by reillumination of the reward tray. The animal had to nose poke and exit the reward tray to start a new trial, with the same two images displayed on the screen. Nose poking the incorrect image (S^-^) resulted in a 5 sec time-out period. The criterion for visual discrimination training was 21/30 correct trials (70% of accuracy) in 60 minutes on two consecutive sessions. Mice were trained for 13 sessions in the Visual Discrimination paradigm. To avoid over-training, mice that reached criterion were rested for two days and then given a reminder trial on the third day. If the mouse performed at least 70% of accuracy on the reminder session, another two days of resting were given. If performance was below 70% of accuracy on the re-train session, the mouse kept training until the criterion of 70% accuracy in two sessions in a row was reached again.

#### Reversal test

Cognitive flexibility can be modeled in rodents through reversal learning paradigms. The correct and incorrect stimuli are switched such that the previously unrewarded stimulus (S^-^) must now be selected (S^+^) to elicit the reward delivery response, and the previously rewarded stimulus (S^+^) is now incorrect (S^-^). Mice were tested in the Reversal task for 16 sessions. The procedure, trials, ITI, time-out periods and criterion of accuracy remained the same as in Visual Discrimination.

### Tissue collection

Twenty-four hours after the last behavioural session, mice were given an overdose of sodium pentobarbital (500 mg/kg, i.p.). Blood samples were collected from the thoracic cavity following a cut to the aorta, and mice were perfused transcardially with 25ml 0.9% saline followed by 25ml 4% paraformaldehyde (Sigma-Aldrich). Blood samples were then centrifuged at 4 °C and 2000×g for 10 min and plasma was stored at −80 °C. Brains were extracted and postfixed in 4% paraformaldehyde overnight, then transferred to 30% sucrose (Fisher Scientific) solution for cryoprotection and remained in the solution until sectioning. Adrenals were extracted and weighed. Brains were sliced into 30-μm coronal sections using a Leica SM2000R microtome (Richmond Hill, ON, Canada). Sections were collected in series of ten throughout the entire rostral-caudal extent of the hippocampus and stored in antifreeze solution consisting of ethylene glycol, glycerol, and 0.1 M PBS at −20°C until immunostaining.

### Doublecortin Immunohistochemistry

Doublecortin (DCX) is a microtubule binding protein expressed in progenitor cells and immature neurons, and is used as a marker for adult neurogenesis (Brown et al., 2003). Briefly, sections were rinsed 3□×□10□min in 0.1□M phosphate buffered saline (PBS), treated with 3% hydrogen peroxide in dH2O for 30□min, rinsed again 3□×□10□min in 0.1□M PBS, and incubated at 4□°C in primary antibody solution: 1:1000, goat anti-doublecortin (Santa Cruz Biotechnology, Santa Cruz, CA, USA). 24□h later, sections were rinsed 5□×□10□min and incubated at 4□°C in an secondary antibody solution with 1:500, rabbit anti-goat (Vector Laboratories, Burlingam, CA,USA). Another 24□h later, sections were rinsed 5 x□10□min and incubated in Avidin-Biotin complex (AB; 1:1000; Vector) for 4□h at room temperature. Sections were then rinsed 3 x 10 min in 0.1M PBS and then washed in 0.175□M sodium acetate buffer 2 times for 2□min, developed using diaminobenzidine in the presence of nickel (DAB Peroxidase Substrate Kit). Sections were then washed 2 x 30 seconds in 0.175□M sodium acetate buffer, rinsed 3 x 10 mins in 0.1M PBS and mounted on slides, and dried. Sections were then dehydrated, cleared with xylene, and coverslipped with Permount (Fisher).

### Microscopy

Microscopy analyses were conducted by an experimenter blind to the group conditions. DCX-expressing cells were quantified in 2 dorsal and 2 ventral sections, using the 40x objective on an Olympus microscope. Cells were classified as DCX-expressing if the stain was dark and cells were present in the inner granule cell layer of the dentate gyrus (Figure 1B-C; see supplemental figure 2 for representative DCX photomicrograph images for each group). Cells were counted separately in the dorsal and ventral regions of the dentate gyrus, as these regions are thought to be functionally distinct. The dorsal region is involved in spatial reference memory whereas the ventral region is involved in working memory, stress and anxiety and has input to the PFC (Fanselow and Dong, 2010; Moser and Moser, 1998; Twining et al., 2020).

Forty cells positively labelled for DCX (20 from the dorsal and 20 from the ventral) were further examined and categorized by morphology. Cells in a proliferative state were classified as type 1 (no or short processes), cells in an intermediate state were classified as type 2 (medium processes with no branching), and cells in a post-mitotic state were classified as type 3 (strong dendritic branching into the molecular layer) (Plümpe et al., 2006; see Figure 1B-C). Two Female Val/Val mice were excluded from these analyses due to tissue loss.

### Plasma Hormone Quantification

Estradiol, progesterone, and dehydroepiandrosterone were measured in plasma samples. A custom steroid hormone panel assay kit (Meso Scale Discovery, Gaitersburg, MD, USA) for all three hormones was used according to manufacturer’s instructions and the inter- and intra-assay coefficients of variation were <10%.

### Vaginal Cytology and Estrus Cycle Assessment

Vaginal lavage samples were taken from female mice on behaviour testing days and estrous stage determined by previous methods (Goldman et al., 2007).

### Statistical Methods

All data were analysed using Statistica software (v. 9, StatSoft, Inc., Tulsa, OK, USA). A general linear analysis of variance (ANOVA) on the total amount of running, body weight, relative adrenal weight, hormone concentration, total average beam breaks and days to reach criteria for the visual discrimination task were calculated with sex (male, female), genotype (BDNF Val/Val, BDNF Met/Met) and exercise (control, AT) as between-group variables. Repeated-measures (ANOVA) with sex (male, female), genotype (BDNF Val/Val, BDNF Met/Met) and exercise (control, AT) as between-group variables, and within-subjects variables of either testing block (1-8), or region (dorsal hippocampus, ventral hippocampus) or DCX expressing cell type (type 1,2,3) were also calculate. Post-hoc comparisons used Newman-Keuls. ANOVA assumptions were tested with Kolmgovox-Smirnov test for normality and Bartlett’s Test of homogeneity of variance, unless otherwise specified these assumptions were not violated. If the assumptions were violated, follow up tests used the Huynh-Feldt correction for repeated measures ANOVA and the Welch’s test for pairwise comparisons. Pearson product-moment correlations were conducted between dependent variables of interest (see supplemental Table 1 for all correlations). A priori we were interested in whether or not sex and genotype influenced the effects of AT on hippocampal neurogenesis and cognitive flexibility. All a priori comparisons used Bonferroni corrections. All effects were considered statistically significant if p ≤ 0.05, trends are discussed if p ≤ 0.10 (Thiese et al., 2016). Outliers were eliminated if higher or lower than 2 standard deviations from the mean. Animals that did not complete 30 trials at least in one session were excluded from the cognitive performance analyses. Data are presented as mean ±standard error and effect sizes for significant results were reported as partial eta squared (η_p_^2^) or Cohen’s d, as appropriate.

## Results

### BDNF Val/Val ran more than BDNF Met/Met mice

BDNF Met/Met ran significantly less than BDNF Val/Val mice regardless of sex (main effect of genotype: [F(1, 29)=13.68 p<0.05, η_p_^2^=0.32; see Table 1; Welch’s t-test p<0.0001). No other significant main or interaction effects were found (all p’s >0.65).

**Table 1.**
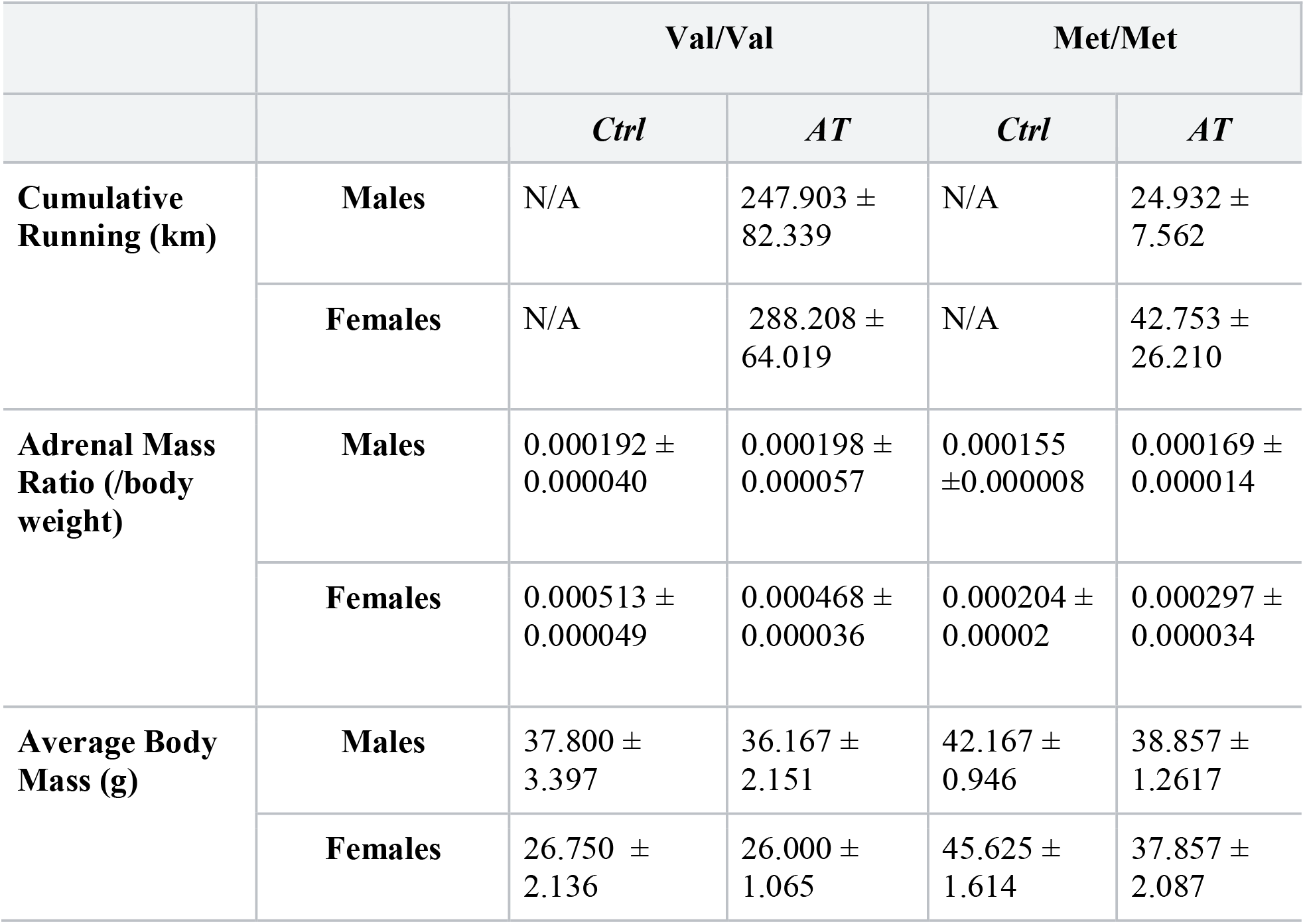
Cumulative running in kilometers (Km), adrenal mass ratio, and body weight across all groups. BDNF Met/Met mice ran significantly less than BDNF Val/Val mice. Adrenal weight expressed as a proportion of body weight. Females had higher relative adrenal weight compared to males, regardless of AT. Female BDNF Met/Met mice had a lower relative adrenal weight than female BDNF Val/Val mice. Female BDNF Val/Val mice had the lowest body weight compared to all other groups. Data presented as mean ± standard error of the mean. Ctrl=Sedentary controls; AT= Aerobic training.

### Relative adrenal mass was higher in females, irrespective of AT

Table 1 shows the adrenal mass expressed in proportion of body weight. Females had larger adrenals in comparison to males regardless of AT (p<0.001; Welch’s t p<0.001; K-W .003). Despite the interaction, females also had larger proportional adrenal mass compared to males in both genotypes (BDNF Val/Val females compared to males (p<0.001, Welch’s p<0.0001; the Met/Met counterparts (p=0.044, Welch’s 0.002, KW p=0.0026; interaction Genotype x Sex (F_1, 41_= 18.2343, p=0.0001, η_p_^2^= 0.308). There were also main effects of genotype (F_1, 41_= 31.710, p<0.001, η_p_^2^= 0.436) and sex (F_1, 41_= 62.458, p<0.001, η_p_^2^=0.604)). There were no other significant main effects or interactions (all p>0.140).

Female BDNF Val/Val mice had significantly lower body mass than female BDNF Met/Met mice as well as both BDNF Val/Val and Met/Met males (Table 1) (genotype by sex interaction [F_1, 41_=19.174, p<0.001, η_p_^2^= 0.318]; post hoc, all p<0.001). There were also main effects of genotype (p<0.001) and sex (p=0.001).

### AT females were more likely to be cycling than sedentary females. BDNF Val/Val mice had higher levels of plasma estradiol and DHEA (males only)

There were more females that were cycling in the AT group than in the control group (p=0.034). BDNF Val/Val mice were also more likely to be regular cycling than BDNF Met/Met mice, although this finding was a trend (p=0.063; see Table 2).

**Table 2.** Vaginal cytology in Female BDNF Val/Val and Met/Met mice. Aerobic training (AT) increased cycling and BDNF Met/Met mice showed lower regular cycling compared to BDNF Val/Val mice. Ctrl=control

|  | Val/Val |  | Met/Met |  |
| --- | --- | --- | --- | --- |
|  | *Ctrl* | *AT* | *Ctrl* | *AT* |
| % Cycling<br>(N/Total N) | 25.0%<br>(2/8) | 62.5%<br>(5/8) | 0.00%<br>(0/8) | 28.6%<br>(2/7) |

We also measured plasma estradiol, progesterone and DHEA. Male BDNF Val/Val mice had higher levels of plasma DHEA than all other groups (p<0.008; F_1, 39_= 4.958, p=0.032, η_p_^2^= 0.112; Figure 2A). There was also a main effect of sex (F_1, 39_= 6.173, p=0.017, η_p_^2^= 0.137) and a trend for a main effect of Genotype (p=0.07) but no other significant main or interaction effects (p<0.118). For plasma estradiol, BDNF Val/Val mice had higher levels of estradiol, regardless of sex or exercise (F_1, 40_= 8.380, p=0.006, η_p_^2^= 0.173; Figure 2B). There were no other significant main or interaction effects of estradiol (all p’s>p=0.375) nor any significant effects seen in plasma progesterone across groups (all p’s >0.118).

**Figure 2.**
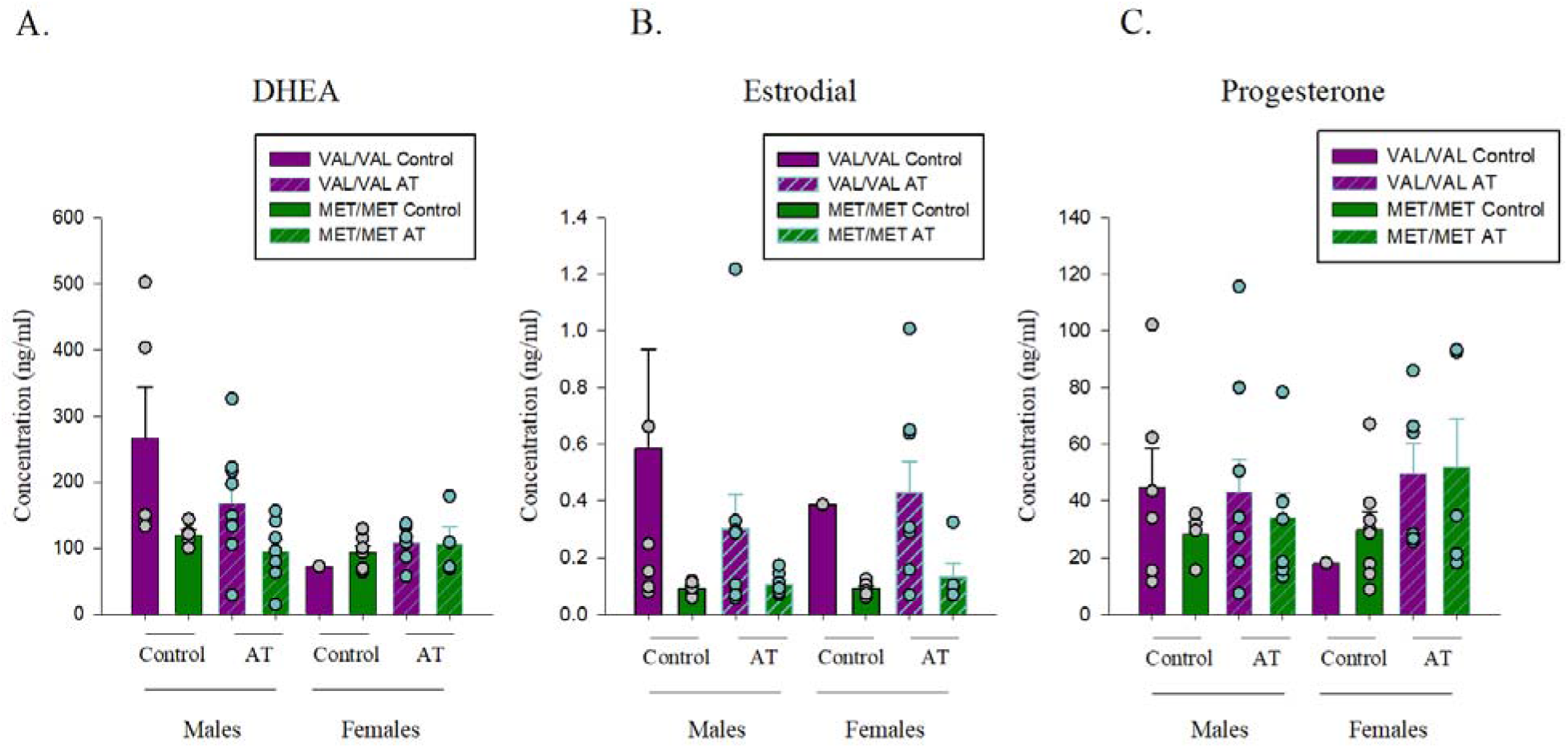
Plasma hormone concentrations (ng/ml) by genotype, sex, and aerobic training (AT) status. A) Dehydroepiandrosterone (DHEA) concentration levels were highest in male BDNF Val/Val mice. B) Estradiol concentration levels were highest in female BDNF Val/Val mice. C) Progesterone concentration levels were not significantly different between groups.

### BDNF Val/Val mice took fewer days to reach criterion on Visual Discrimination trials and AT reduced days to reach criterion in BDNF Met/Met mice only

Figure 3A shows the days to reach training criterion for visual discrimination. The analysis revealed that sedentary control BDNF Met/Met mice reached the training criterion in more days than control BDNF Val/Val (p<0.001, Cohen’s d= 1.909; interaction of genotype by AT (F_1,56_= 13.333, p<0.001, η_p_^2^= 0.19, Welch’s t=p<0.0001)). However, BDNF Met/Met mice showed a reduction in days to reach criteria with AT (p=0.003, Cohen’s d=1.269; Welch’s t: p=0.003) but the same AT benefit was not statistically significant in BDNF Val/Val mice (p=0.229; Welch’s t: p=0.136). There were also main effects of genotype (F_1,56_= 48.533, p<0.001, η_p_^2^= 0.46) and AT(F_1,56_=4.082, p=0.048, η_p_^2^= 0.07) but no other significant main effects or interaction effects were observed.

**Figure 3.**
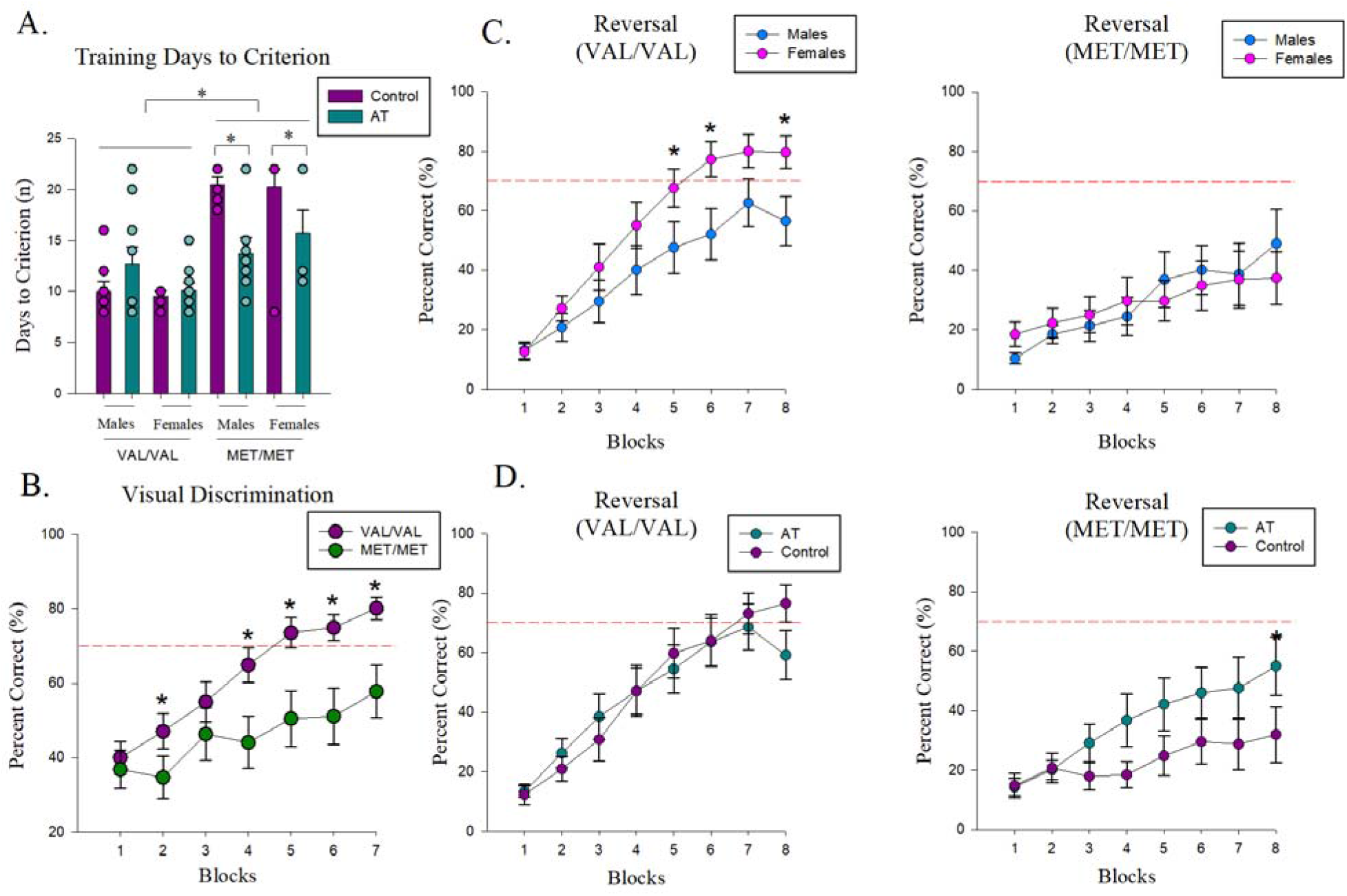
A) BDNF Met/Met mice took more days to reach training criterion than BDNF Val/Val mice, regardless of sex. Aerobic training (AT) reduced days to reach criterion in BDNF Met/Met mice only. (*p<0001 vs Met/Met sedentary control; individual data points are plotted but due to a number of mice achieving criterion on the same day there are many overlapping symbols, the sample sizes are male Val/Val control n=8, male Val/Val AT n= 10, female Val/Val controls n= 8, female Val/Val AT n=8, male Met/Met control n=6, male Met/Met AT n= 7, female Met/Met controls n=8, female Met/Met AT n=7). B)Visual discrimination training. BDNF Met/Met mice had fewer correct choices when learning to discriminate the visual stimulus compared to BDNF Val/Val mice. C-D) Reversal training (cognitive flexibility). C) Female BDNF Val/Val mice had better cognitive flexibility, regardless of AT, than males of the same genotype (on blocks 5, 6 and 8 (all p’s<0.05; genotype by sex by block (F_7,301_= 2.259, p=0.030, η_p_^2^= 0.050). No differences were observed in BDNF Met/Met mice (p’s >0.610). D) Although Met/Met mice did not reach criterion for visual discrimination or reversal, AT increased cognitive flexibility (reversal performance) in BDNF Met/Met mice compared to sedentary controls (block 8 p=0.025; genotype by AT by block (F_7,301_= 2.287, p=0.028, η_p_^2^= 0.050). No significant differences were observed in BDNF Val/Val mice (all p’s>0.14). Red dotted lines indicate the criterion of 70% correct trials.

Across the blocks of visual discrimination trials, BDNF Met/Met mice had less correct responses than BDNF Val/Val mice (on blocks 2, 4-7 (all p’s <0.02; genotype by block: F_6,258_= 4.139, p=0.0005 (Huynh-Feldt correction: 0.002254), η_p_^2^= 0.087; Figure 3B). A priori, we expected AT mice to perform better than control mice and they did on the fourth and last block of sessions (p’s <0.0007; AT by block: F_6,258_= 1.849, p=0.09, η_p_^2^= 0.041). There were also main effects of blocks (F_6,258_= 33.409, p<0.0001(Huynh-Feldt correction: p <0.0001), η_p_^2^= 0.437) and genotype (F_1,43_= 5.138, p=0.028, η_p_^2^= 0.107) but no other significant effects (all p’s >0.23).

Given the differences in overall running between the genotypes we also examined activity levels in the touch screen chambers. Examining the average number of beam breaks across all sessions of either visual discrimination acquisition or reversal there were no significant main or interaction effects, except for a strong trend for females to show more activity than males in acquisition (F_1,56_= 3.75, p=0.058, η_p_^2^= 0.063) but not under reversal trials (all p’s>0.18; see Table 3).

**Table 3.** Average beam breaks across all sessions in visual discrimination acquisition and reversal stages. Females tended to have more activity than males in visual discrimination acquisition regardless of genotype (p=0.063). Data presented as mean ± standard error of the mean. Ctrl=control, AT=aerobic training

|  |  | Val/Val |  | Met/Met |  |
| --- | --- | --- | --- | --- | --- |
|  |  | *Ctrl* | *AT* | *Ctrl* | *AT* |
| **Acquisition** | **Males** | 558.202±<br>57.661 | 581.267±<br>50.538 | 701.624±<br>82.949 | 668.286±<br>54.929 |
|  | **Females** | 807.962±<br>158.275 | 757.016±<br>77.982 | 784.528±<br>135.467 | 718.798±<br>118.221 |
| **Reversal** | **Males** | 599.919±<br>68.200 | 613.508±<br>73.906 | 721.229±<br>87.315 | 701.268±<br>57.066 |
|  |  | *Ctrl* | *AT* | *Ctrl* | *AT* |
|  | **Females** | 770.653±<br>139.041 | 762.153±<br>73.299 | 669.617±<br>115.676 | 691.286±<br>132.881 |

### Female BDNF Val/Val showed increased cognitive flexibility compared to males, regardless of AT. AT improved cognitive flexibility in Met/Met mice regardless of sex

Female BDNF Val/Val mice had better cognitive flexibility (reversal performance), regardless of AT, than males of the same genotype (on blocks 5, 6 and 8 (all p’s<0.05; genotype by sex by block (F_7,301_= 2.259, p=0.030, η_p_^2^= 0.050; Huynh-Feldt correction: p =0.040) with no significant differences between the sexes of BDNF Met/Met mice (all p’s >0.610). In addition, the BDNF Met/Met mice had fewer correct responses on reversal learning than BDNF Val/Val mice, moreso in females (Block 4-8 all p’s <0.012) compared to males (Block 7, p=0.016). In addition, BDNF Met/Met mice had fewer correct responses compared to BDNF Val/Val mice on the reversal task particularly under sedentary control conditions (Blocks 4-8, p’s <0.002) than under AT (Block 7 p=0.040). Although Met/Met mice did not reach criterion for visual discrimination or reversal, better cognitive flexibility (reversal performance) was observed in AT BDNF Met/Met mice compared to sedentary control (block 8 p=0.025; genotype by AT by block (F_7,301_= 2.287, p=0.028, η_p_^2^= 0.050; Figure 3D; Huynh-Feldt correction: p =0.024). AT did not significantly influence reversal training in BDNF Val/Val mice across blocks (all p’s>0.14). There were also significant main effects of genotype (F_1,43_= 9.193, p=0.004, η_p_^2^= 0.176) and block (F_7,301_= 43.043, p<0.001, η_p_^2^= 0.500) and an interaction of blocks by genotype (F_7,301_= 6.190, p<0.0001, η_p_^2^= 0.125) but no other significant effects (all p’s>0.210).

### Voluntary running increased the total density of doublecortin-expressing cells in the ventral region of BDNF Val/Val mice

AT increased the total density of DCX-expressing cells in the ventral region compared with sedentary controls in the BDNF Val/Val mice (p=0.0097; Cohen’s d= 0.393; Welch’s: p=0.04) but not in the BDNF Met/Met mice (p=0.935; interaction between genotype, AT, and hippocampal region [F(1,53)=4.373, p=0.041, η_p_^2^=0.076; see Figure 4A-B]. Further a trend was found for an interaction of genotype by sex by region [F(1,53)=3.976, p=0.051, η_p_^2^=0.069], but no meaningful post-hoc significant outcomes were found, and a main effect of hippocampal region was found [F(1,53)=39.958, p<0.0001, η_p_^2^=0.430]). In addition, a trend for a main effect of AT [with the AT group having more DCX-expressing cells: F(1,53)=3.454, p=0.069, η_p_^2^=0.061]) was found but no other significant main or interaction effects (p’s >0.34).

**Figure 4.**
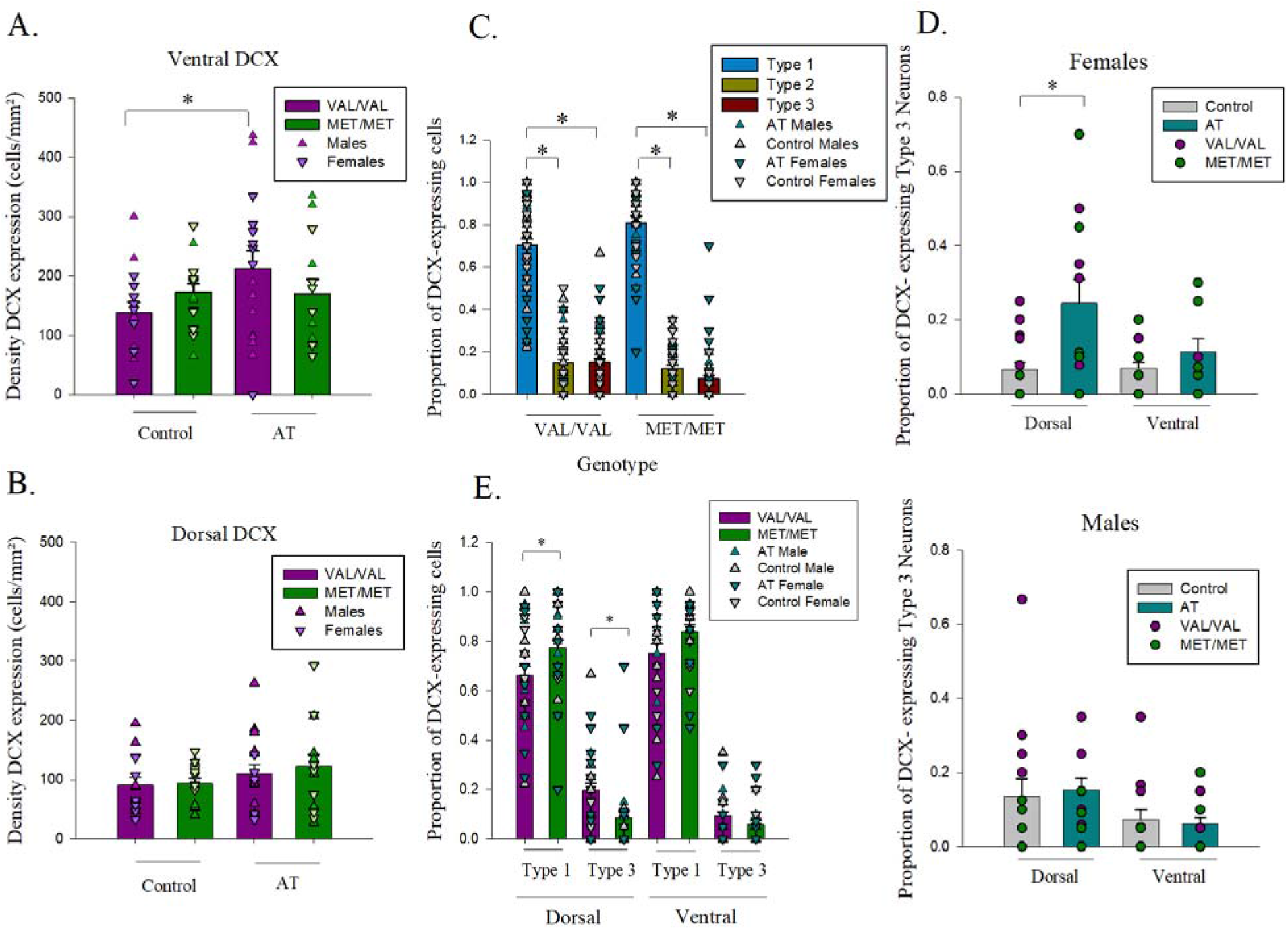
A-B) Density of doublecortin (DCX)-expressing cells by BDNF genotype and aerobic training (AT) in the ventral (A) and dorsal (B) region of the dentate gyrus. The density of doublecortin-expressing cells was increased with AT in the ventral (A), but not dorsal (B), region of the hippocampus in the BDNF Val/Val mice. C-E) Maturity of doublecortin (DCX) expressing cells. C) There was a significantly greater proportion of proliferative type 1 DCX expressing cells compared to both type 2 and type 3 in both the BDNF Val/Val and Met/Met mice. D) The proportion of type 3 DCX-expressing cells increased with AT in female but not male mice. E) BDNF Met/Met mice had a greater proportion of type 1 DCX-expressing cells in the dorsal region of the hippocampus compared to the BDNF Val/Val mice. Whereas, the BDNF Val/Val mice had a greater proportion of type 3 DCX-expressing cells compared to the Met/Met mice, in the dorsal hippocampus. See supplemental Figure 1 for graphs separated by all groups.

### Voluntary running increased the proportion of Type 3 doublecortin-expressing cells in the dorsal region of females; Val/Val mice had more type 3 doublecortin-expressing cells

The morphology of the DCX-expressing cells indicates the maturity of the new neurons (Plümpe et al., 2006) and we next examined the proportion of type 1, 2 and 3 DCX-expressing cells. Type 1 DCX-expressing cells were significantly higher in the ventral compared to the dorsal region, and conversely, type 3 DCX-expressing cells were significantly higher in the dorsal compared to the ventral region (post-hocs, all p’s<0.05; a region by type of DCX-expressing cell interaction (F(2,104)=8.2627 p=0.0005, η_p_^2^=0.137]; Huynh-Feldt correction: p=<0.0001). There was a significantly greater proportion of type 1 DCX-expressing cells compared to both type 2 or 3 in both BDNF Val/Val and Met/Met mice (all p’s<0.0001; type by genotype interaction (F(2,104)=329.675 p<0.0001, η_p_^2^=0.8638]; Huynh-Feldt correction: p =0.031) (see Figure 4C). Despite the genotype by sex by group interaction effect ([F(1,52)=39219 p<0.0001, η_p_^2^=0.999]; Huynh-Feldt correction: p =0.0125]), there were no significant differences between the groups of interest in the post hoc analysis.

Type 3 DCX-expressing cells were increased in the AT group compared to sedentary control in female, but not male mice (a priori p=0.001, Cohen’s d=0.2491; Figure 4D Welch’s t: p=0.018). Furthermore, proportional maturity varied by genotype in the dorsal region only. Met/Met mice had a greater proportion of type 1 cells compared to the Val/Val mice (p=0.003, Cohen’s d=0.626; Figure 4C), whereas the opposite was true for the dorsal type 3 DCX expressing cells where Met/Met mice had significantly less type 3 DCX expressing cells compared to the Val/Val mice (p=0.003, Cohen’s d=0.752; Figure 4E).

### Plasma dehydroepiandrosterone (DHEA) was related to cognitive performance, dependent on genotype and sex

To understand how hormones and other outcome variables may be related we performed correlations. DHEA was positively related to average cognitive flexibility in female BDNF Val/Val mice (r=0.937, p=0.006) and positively related to visual discrimination in male BDNF Met/Met mice (r=0.767, p=0.004). Plasma progesterone was negatively related to visual discrimination in female BDNF Met/met mice only (r=−0.796, p=0.001) There were no other significant correlations by sex or genotype on any of the dependent measures that survived Bonferroni correction. All correlations are shown in Supplemental Table 1.

## Discussion

BDNF Met/Met mice had fewer correct responses on the visual discrimination task compared to BDNF Val/Val mice regardless of sex or AT. Female BDNF Val/Val mice performed better than males on the cognitive flexibility task, regardless of AT. AT benefited both cognitive flexibility and visual discrimination in BDNF Met/Met mice regardless of sex. AT significantly increased neurogenesis in the ventral region of the hippocampus in middle-aged BDNF Val/Val mice (wild-type), but not in the mice with the BDNF Val66Met polymorphism (BDNF Met/Met mice), regardless of sex. However, when we examined the specific effect of voluntary AT on the maturity of these DCX-expressing cells, AT increased the proportion of type 3 DCX-expressing cells in the dorsal region in female mice compared to males, regardless of genotype.

Furthermore, BDNF Met/Met mice were found to have a lower proportion of dorsal type 3 DCX-expressing cells, along with fewer correct responses during visual discrimination, compared to the BDNF Val/Val mice. In summary, our results indicate that in middle-age, female mice may benefit more in terms of neuroplasticity and cognitive flexibility with AT, an effect that can be influenced by the BDNF Val66Met polymorphism. Further we find that BDNF Met/Met mice showed cognitive benefit from AT, even though they did not run to the same extent as BDNF Val/Val mice, indicating that there may be a lower threshold for AT activity to influence cognition based on genotype. Further, in our study we find divergent effects of AT that depend on both genotype and sex in middle-aged mice.

### Cognitive flexibility performance in middle-age is associated with sex and genotype

Here we found that BDNF Met/Met middle-aged mice made fewer correct choices on both visual discrimination and cognitive flexibility than BDNF Val/Val middle-aged mice, regardless of sex or AT. Previous studies have found in middle-age humans that individuals with the BDNF Met/Met genotype show a steeper decline in executive function and episodic memory (Boots et al., 2017), but others have found less decline in cognitive flexibility assessed through set-shifting in patients with Parkinson’s Disease (van der Kolk et al., 2015) or in healthy aging using the Digital Symbol Substitution Test (Barha et al., 2019b). The inconsistencies among studies may be due to tasks given (set shifting, processing speed, flexibility), health status (Parkinson’s versus cognitive healthy aging), and/or relative proportions of males to females. Indeed, in our study we found that female BDNF Val/Val mice showed greater cognitive flexibility than males regardless of AT. These findings are at least partially consistent with previous literature where human females had a higher cognitive baseline compared to males in middle-age for executive functions (Levine et al., 2021) and where white female individuals with the BDNF Val/Val genotype had better digit symbol substitution scores than white males (Barha et al., 2019b). These findings suggest that the female sex is associated with better cognitive flexibility in both humans and rodents, that may depend on genotype. The sex by genotype differences in cognitive flexibility may be partially modulated by differences in gene expression in the CA3 region (Marrocco et al., 2017), as the CA3 region, along with the dentate gyrus contributes to cognitive flexibility (Berdugo-Vega et al., 2021; Webler et al., 2019).

### AT improved cognitive performance in BDNF Met/Met mice but increased neurogenesis in BDNF Val/Val mice

In our study middle-aged BDNF Met/Met mice did not run as much as middle-aged BDNF Val/Val mice, regardless of sex. Previous reports have shown no differences in the amount of running between BDNF Val/Val and BDNF Met/Met in young adult mice (Sandrini et al., 2019; Ieraci et al., 2016), therefore, it is possible that age influences the running pattern in BDNF Met/Met. The increased amount of running by the BDNF Val/Val groups did not translate into greater mobility during the training in the touchscreen chambers as there were no activity differences between genotypes during visual discrimination or reversal training. In addition, although there were differences between genotypes in body weight in the females, there were no significant correlations between body weight and cumulative running (see supplemental Table 1). It is difficult to know why the BDNF Met/Met mice ran less than the BDNF Val/Val mice but future studies should explore different ways to increase the amount of running in these mice, perhaps using forced exercise paradigms (see below).

Despite the fact that BDNF Met/Met mice did not run as much as BDNF Val/Val mice, we found greater AT efficacy to promote both visual discrimination and cognitive flexibility in the BDNF Met/Met mice, regardless of sex. However, it is important to note that despite the AT-induced improvements in BDNF Met/Met mice, they still performed worse than BDNF Val/Val mice. Intriguingly, individuals with the BDNF Met/Met genotype have a slower rate of cognitive decline (Barha et al., 2019b, van der Kolk et al., 2015). It is possible that given exercise is an effective strategy to ward off cognitive decline, perhaps exercise benefits BDNF Met/Met individuals more than others. Thus, it may be that the slower rates of decline in executive functions seen in Met carriers, may be related to the greater cognitive benefits of AT in Met carriers. Other studies have found that AT enhances cognition in older human females compared to older males (Liu-Ambrose et al., 2018), and although we did not find a sex by genotype difference in these middle-aged mice, it is possible that our mice were too young to see this aging effect. A previous meta-analysis reported greater AT benefits on non-spatial memory tasks in healthy male older rodents compared to female rodents (Barha et al., 2017b). Similarly, Short et al., (2022) found beneficial effects of AT on cognitive flexibility in male, but not in female, middle-aged mice. It is possible that different genotypes and sexes require different amounts of AT to improve cognition, and future studies could explore longer AT interventions in the Val/Val mice.

BDNF Val/Val mice showed an AT-induced increase in ventral DCX-expressing cells compared to sedentary controls, that was not seen in BDNF Met/Met mice. It is tempting to speculate that this was due to the fact that the BDNF Met/Met mice did not run as much as the BDNF Val/Val mice. However, the lack of an AT effect on neurogenesis in BDNF Met/Met is consistent with previous research done in younger male mice (Ieraci et al., 2016), even though these younger male mice did run as much as the younger BDNF Val/Val mice (Ieraci et al., 2016). Taken together with our results, genotype seems to play a greater role, independent of the amount of running, to thwart the AT-induced increase in neurogenesis. A possible explanation for this genotype effect is the impaired release, binding affinity and intracellular trafficking of neural BDNF that is seen with the Val66Met polymorphism (Chen et al., 2008; Chiaruttini et al., 2009). As BDNF increases the survival of new neurons and thus increases neurogenesis (Sairanen et al., 2005), less BDNF secretion (due to lack of AT or impairments intrinsic to the genotype) could reduce the amount of neurogenesis in the BDNF Met/Met mice. Indeed, crucial BDNF transcripts for dendritic trafficking and optimal neuroplasticity are differentially expressed in the hippocampus of BDNF Met/Met and Val/Val young mice (Mallei et al., 2015) and future studies should confirm if similar effects occur in middle-aged rodents. Our findings that AT improved cognitive flexibility in BDNF Met/Met mice but increased neurogenesis in BDNF Val/Val mice, also suggest that AT promotes neuroplasticity and cognition based on divergent pathways that are expressed differently by genotype.

### Female mice showed more mature DCX-expressing cells than male mice with AT

In understanding the role that neurogenesis may play the ability of AT to promote cognition and reduce cognitive decline with aging, it is important to consider the maturity of the new neurons. AT increased the proportion of dorsal type 3 DCX-expressing cells, an effect that was driven by the female mice, regardless of genotype, agreeing with previous literature where AT increases neurogenesis (van Praag et al., 1999), though sex differences were not examined in that study. As the dorsal hippocampus is involved in memory (Fanselow and Dong, 2010), and AT exercise provides cognitive benefits in a variety of species (Barha et al., 2017b; Erickson et al., 2011; Makizako et al., 2015; ten Brinke et al., 2015), increased neurogenesis in this region may be expected. Type 3 DCX-expressing cells are classified as late progenitor cells and are mature DCX-expressing cells (Plümpe et al., 2006). Intriguingly, the running-induced increase in neurogenesis was driven by female mice, consistent with previous results indicating that voluntary or treadmill running increased neurogenesis to a greater degree in younger adult female mice compared to males (Ma et al., 2012; Ransome and Hannan, 2013). One caveat to the past findings is that typically younger female rodents run more than male rodents (Ransome and Hannan, 2013), but in our sample of middle-aged C57B6 mice, there was no sex difference in the total amount of running, suggesting that despite similar levels of AT, a greater neurogenic response was seen in middle-aged female mice compared to middle-aged male mice. Our finding that neurogenesis was increased in AT female mice is consistent with our behaviour results wherein the females outperformed the males on both cognitive tasks. Future studies are needed to further investigate these sex differences found in response to AT in middle-age.

A lower proportion of type 3 DCX-expressing cells was also found to be present in the dorsal region of the hippocampus of BDNF Met/Met compared to the BDNF Val/Val mice. As mentioned above, type 3 DCX-expressing cells are mature or late progenitor cells (Plümpe et al., 2006), therefore BDNF Met/Met mice contain a lower proportion of mature DCX-expressing cells compared to Val/Val mice. This may be due to the lower neural release of BDNF in BDNF Met/Met mice compared to the BDNF Val/Val mice. Intriguingly, DCX-expressing cells are more likely to be activated with cognitive flexibility in male rats (Webler et al., 2019), suggesting that this increase may have contributed to the better flexibility in the female BDNF Val/Val mice in the present study. Another intriguing finding is that Met/Met mice had more type 1 DCX-expressing neurons compared to Val/Val mice, with the reverse for Type 3 DCX-expressing cells. This suggests that the BDNF Met/Met mice have fewer new neurons surviving to maturity, or have slower maturation, as Met/Met individuals have impaired BDNF/TrKB signaling that strongly influences the survival and maturation of hippocampal neurons (Donovan et al., 2008). Given that one study has noted that maturation of DCX-expressing cells may be differentially involved in cognitive flexibility (Webler et al., 2019), and that in our studies female BDNF Val/Val mice had superior cognitive flexibility and greater proportion of mature DCX cells these findings may be linked. The contribution of different ages of new neurons to cognitive flexibility across sex and genotype is an important route for future studies.

The BDNF Val66Met polymorphism is associated with reduced neurological functioning such as impaired hippocampal-dependent memory (Chen et al., 2008; Dincheva et al., 2014; Egan et al., 2003; Kennedy et al., 2015; Lamb et al., 2015) and compromised PFC activity (Pattwell et al., 2012). One question that arises is why is the BDNF Met/Met SNP conserved given its prevalence in neurological functioning and disorders? Here, we found that BDNF Met/Met mice had greater AT efficacy to promote both visual discrimination and cognitive flexibility, regardless of sex. This along with other potential advantages of the BDNF Met/Met SNP such as slower rates of cognitive decline (Barha et al., 2019b, van der Kolk et al., 2015), and protection against cancer-related fatigue (Feng et al., 2020), may in part explain why this SNP is conserved in many populations.

### Neurogenesis does not always mirror cognitive performance

AT significantly increased neurogenesis in the ventral region of the hippocampus in middle-aged BDNF Val/Val mice (wild-type), but not in the mice with the BDNF Val66Met polymorphism (BDNF Met/Met mice), regardless of sex. However, when we examined the specific effect of voluntary AT on the maturity of these DCX-expressing cells, AT increased the proportion of type 3 DCX-expressing cells in the dorsal region in females compared to males, regardless of genotype. Furthermore, BDNF Met/Met mice were found to have a lower proportion of dorsal type 3 DCX-expressing cells, along with impaired visual discrimination, compared to the BDNF Val/Val mice. This latter finding suggests that a more fruitful way to find coherence between neurogenesis measures and cognition is to look at changes in maturation of new neurons with performance as different stages of neurogenesis are more likely to contribute to different forms of memory, including cognitive flexibility (Deng et al., 2010; Nakashiba et al., 2012; Webler et al., 2019; Yagi et al., 2022).

It is tempting to equate enhanced neurogenesis with improved performance on cognitive tasks, but this relationship is not always seen in the literature. Seizures and traumatic brain injury (TBI) increase neurogenesis in the hippocampus but neither of these are associated with improved cognition (Cho et al., 2015; Jessberger et al., 2007). Indeed, studies have found that TBI or seizure-enhanced neurogenesis contributed directly to the cognitive deficits (Cho et al., 2015; Jessberger et al., 2007). It is not just the number of new neurons that are produced but whether or not they are active and contributing positively to the function of the hippocampus. There is likely an optimal level of neurogenesis that is needed for optimal performance that may differ across genotype and sex (Lambert et al., 2019).

### Plasma DHEA was related positively to cognitive performance in middle-aged mice and both estradiol and DHEA were higher in BDNF Val/Val mice

Although both males and females show increased cortisol levels in older age, this effect is three times more pronounced in females compared to males (Otte et al., 2005). Furthermore, increased cortisol levels in older age are linked to poorer cognition and smaller hippocampal and prefrontal cortical volumes (Stomby et al., 2016). Additionally, individuals carrying the Met allele show reduced cortisol release in response to stress, particularly in females (Fiocco et al., 2020; Jiang et al., 2017) compared to individuals with the BDNF Val/Val genotype. This effect is mimicked in male BDNF Met/Met mice with increased corticosterone and adrenocorticotropic hormone after 7 days of restraint stress (Yu et al., 2012). However, in the present study we did not observe any significant differences in relative adrenal mass with AT. Thus, future studies could further examine this effect and additionally measure corticosterone level differences in response to AT and the BDNF Val66Met polymorphism.

Furthermore, exercise may benefit females by extending their cyclicity as in the present study we found that more females were cycling in the exercise group compared to the control group, regardless of genotype. Estrous cycle characteristics in younger female mice did not differ by BDNF genotype, although there were effects of genotype by estrous cycle phase on anxiety and hippocampus-dependent memory (Spencer et al., 2010; Bath et al 2012). However, to our knowledge no study has examined the estrous cycle in middle-aged mice by BDNF genotype. In our study we also found that estradiol was higher in BDNF Val/Val mice which mirrors our findings of more regular cycling in the BDNF Val/Val mice compared to BDNF Met/Met mice. Intriguingly, hippocampal activation with working memory is differentially activated with estradiol in young adult women based on BDNF genotype (Wei et al., 2018), indicating that estradiol may have different functional implications for cognition based on genotype. Future studies should explore how estrogens may mediate cognitive flexibility in middle-aged mice across the BDNF genotype.

In the present study, DHEA levels were increased in male BDNF Val/Val middle-aged mice and in both sexes plasma DHEA levels were related positively to cognition. In humans, exercise was associated with improved cognition and DHEA levels in older women (Farzane and Koushkie Jahromi, 2022), partially consistent with these findings. Another intriguing area of research is the myokine, irisin, which is increased with exercise, mediates the BDNF-induced increase with exercise (Jodeiri Farshbaf and Alvina, 2021) and interferes with ovarian growth and fertility (Luo et al., 2021), suggesting that the FNDC5/irisin pathway may be a fruitful avenue to explore. In the present study, although we did not see large effects of AT on hormones this may be due to the age of the mice or the amount of physical activity.

### Future directions and Limitations

The BDNF Met/Met mice in our study did not run as much as the BDNF Val/Val mice and performed worse on both cognitive tasks. In the future, one strategy might be to start cognitive training earlier or extend it longer (to give more time to reach training criteria) and take measures to ensure more running in the BDNF Met/Met. This also might suggest that interventions in BDNF Met/Met individuals need to start earlier in one’s lifespan. Previous studies found that after AT, BDNF levels increased more in females compared to males (Barha et al., 2017a, 2017b, 2017c). However, as we did not measure hippocampal BDNF levels, we cannot positively link our results of improved female cognition and neurogenesis to higher BDNF levels compared to males and future studies should be encouraged to examine BDNF levels.

In the future, given that this study saw differences in the amount of cumulative running, researchers should consider a way to titrate the amount of running using paradigms involving restricted access or forced exercise. However, the use of shocks or punishment to ensure movement on the forced treadmills will increase stress which will also affect cognition and neurogenesis and will thus affect interpretation of the findings.

The PFC is important for cognitive flexibility and should be studied in the future. Given that the PFC is not normally neurogenic, other indicators for neuroplasticity could be evaluated, such as synaptophysin, synaptic structural complexity and synapses number, given that physical exercise benefits those markers in the PFC (Mu et al., 2022).

The findings that AT increased neurogenesis in the ventral hippocampus is interesting because it is the region that has more connections with the PFC and plays a pivotal role in inhibitory processes and emotional regulation (Adhikari et al., 2010; Chen et al., 2018). Also, although physical exercise increases the number of new neurons in the hippocampus (Gualtieri et al., 2017); Connolly et al., 2022), the effects on depressive and anxious-like behavior in middle-aged subjects are ambiguous (Morgan et al., 2018; Chaiton et al., 2019). Indeed, Chaiton et al. (2019) used female middle-aged mice and found that increased neurogenesis was not linked to changes in depressive-like behaviour. BDNF Met/Met individuals have a depressive phenotype (Notaras et al., 2017) that could perhaps benefit from exercise. Future studies should consider the effects of physical activity on depressive-like behaviour.

### Conclusions

Female mice with the wild-type BDNF Val/Val alleles showed greater cognitive flexibility regardless of AT, compared to all other groups. AT increased neurogenesis and improved cognitive measures differentially depending on genotype, as AT improved visual discrimination and cognitive flexibility in the BDNF Met/Met mice, but increased neurogenesis in the ventral region of BDNF Val/Val mice. AT also increased the proportion of mature DCX expressing cells in the dorsal region of the hippocampus in female mice, regardless of genotype. Thus, AT may exert its effects on neuroplasticity and cognition differentially by genotype. This study also indicates that middle-aged female mice may benefit more from AT in terms of improved cognitive flexibility and neuroplasticity, an effect that can be influenced by the BDNF Val66Met polymorphism.

## Supporting information

Supplemental Section

## Acknowledgements

This research was funded by a Canadian Institutes of Health Research grant (SVB-145582) awarded to CKB, TLA and LAMG. DIP was funded by Consejo Nacional de Ciencia y Tecnología, México (Postdoctoral grant, I-P D. 327002).

We thank Francis Lee for permission to use the mice and Matt Hill for supplying the original breeding pairs.

## Supplemental Section

**Supplemental Figure 1.**
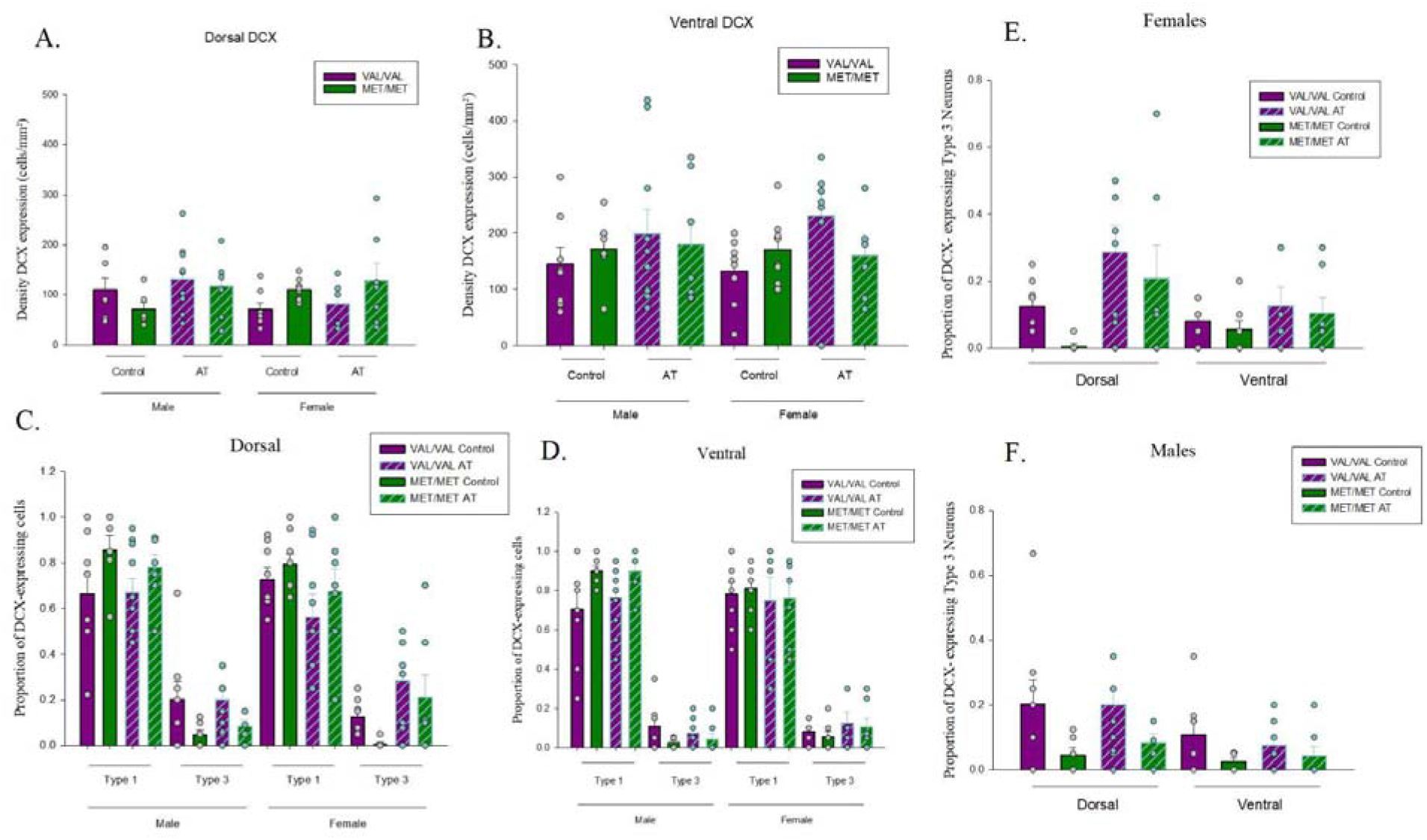
Main findings separated by all groups. Dots represent individual data points. A-B) Density (cells/mm2) of doublecortin (DCX)-expressing cells by BDNF genotype, sex, and aerobic training (AT) in the dorsal (A) and ventral (B) region of the dentate gyrus. C-D) Maturity of DCX-expressing cells (type 1, and type 3) by BDNF genotype, sex, and AT in the dorsal (C) and the ventral (D) region of the dentate gyrus. E-F) The proportion of mature type 3 DCX-expressing cells by BDNF genotype, region, and AT in female (E) and male (F) mice.

**Supplemental Figure 2.**
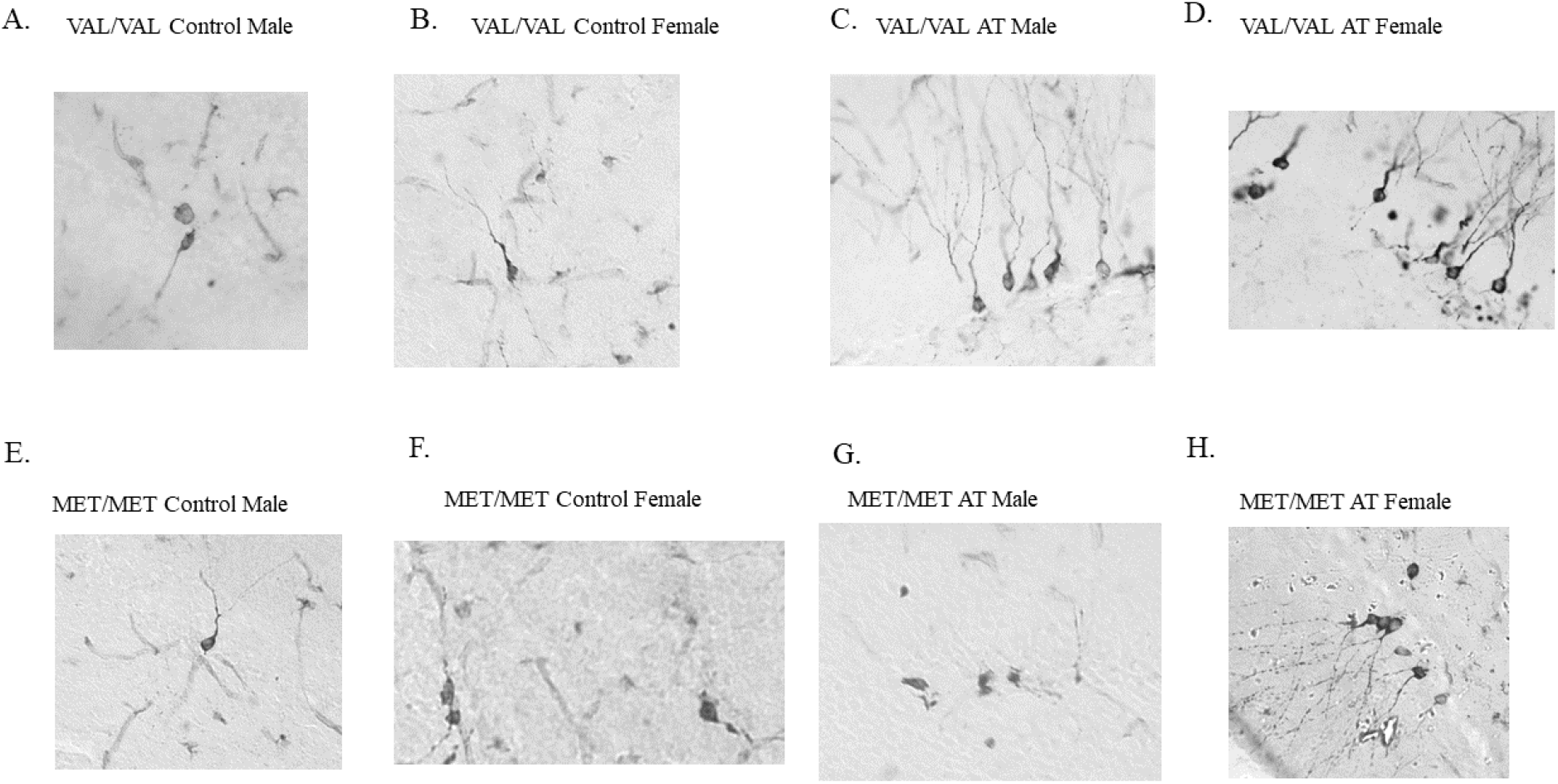
Photomicrographs of doublecortin (DCX) expressing cells at 40x objective. A-D) BDNF VAL/VAL mice. E-H. BDNF MET/MET mice. A) Photomicrograph of a BDNF VAL/VAL control male mouse. B) Photomicrograph of a BDNF VAL/VAL control female mouse. C) Photomicrograph of a BDNF VAL/VAL aerobic training (AT) male mouse. D) Photomicrograph of a BDNF VAL/VAL AT female mouse. E) Photomicrograph of a BDNF MET/MET control male. F) Photomicrograph of a BDNF MET/MET control female. G) Photomicrograph of a BDNF MET/MET AT male. H) Photomicrograph of a BDNF MET/MET AT female.

**Supplemental Figure 3.**
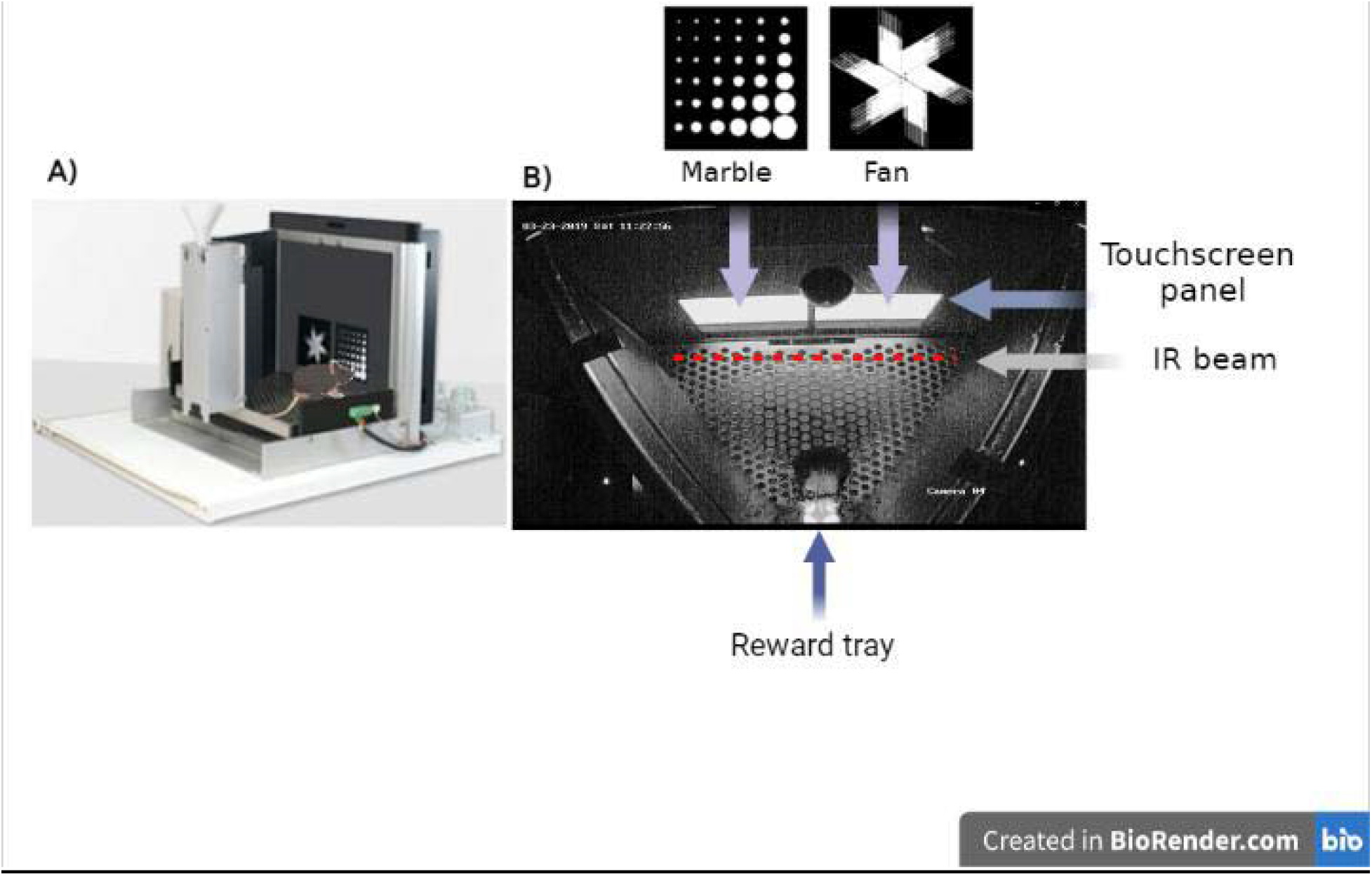
A) Illustrated side view of the touchscreen chamber (modified from: edspace.american.edu/openbehavior/project/touchscreen-cognition-mousebytes/). Panel B) shows a real image from above of the trapezoidal chamber and the touchscreen panel that displays the stimulus (upper) and the reward tray (bottom).

**Supplemental Table 1.**
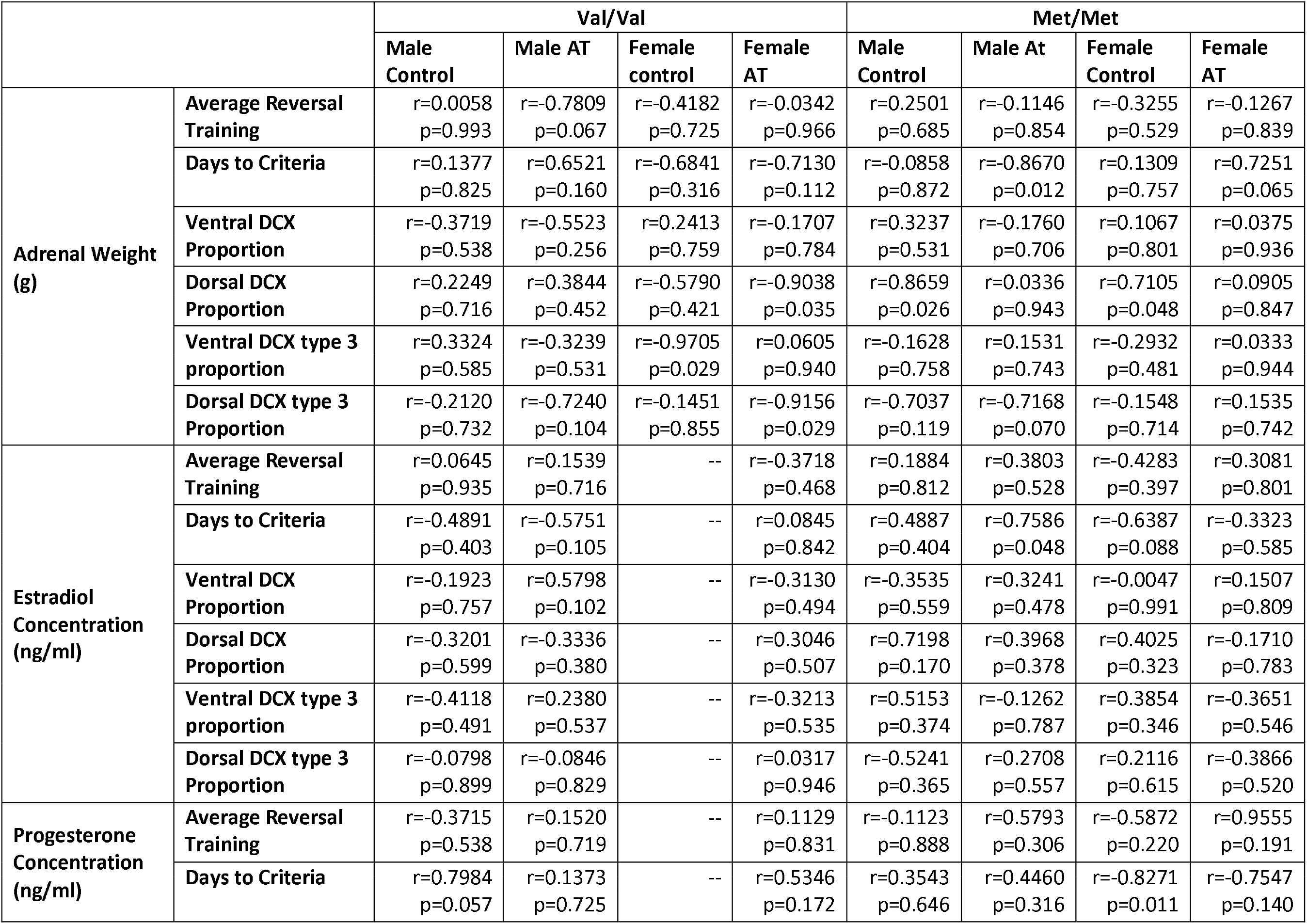
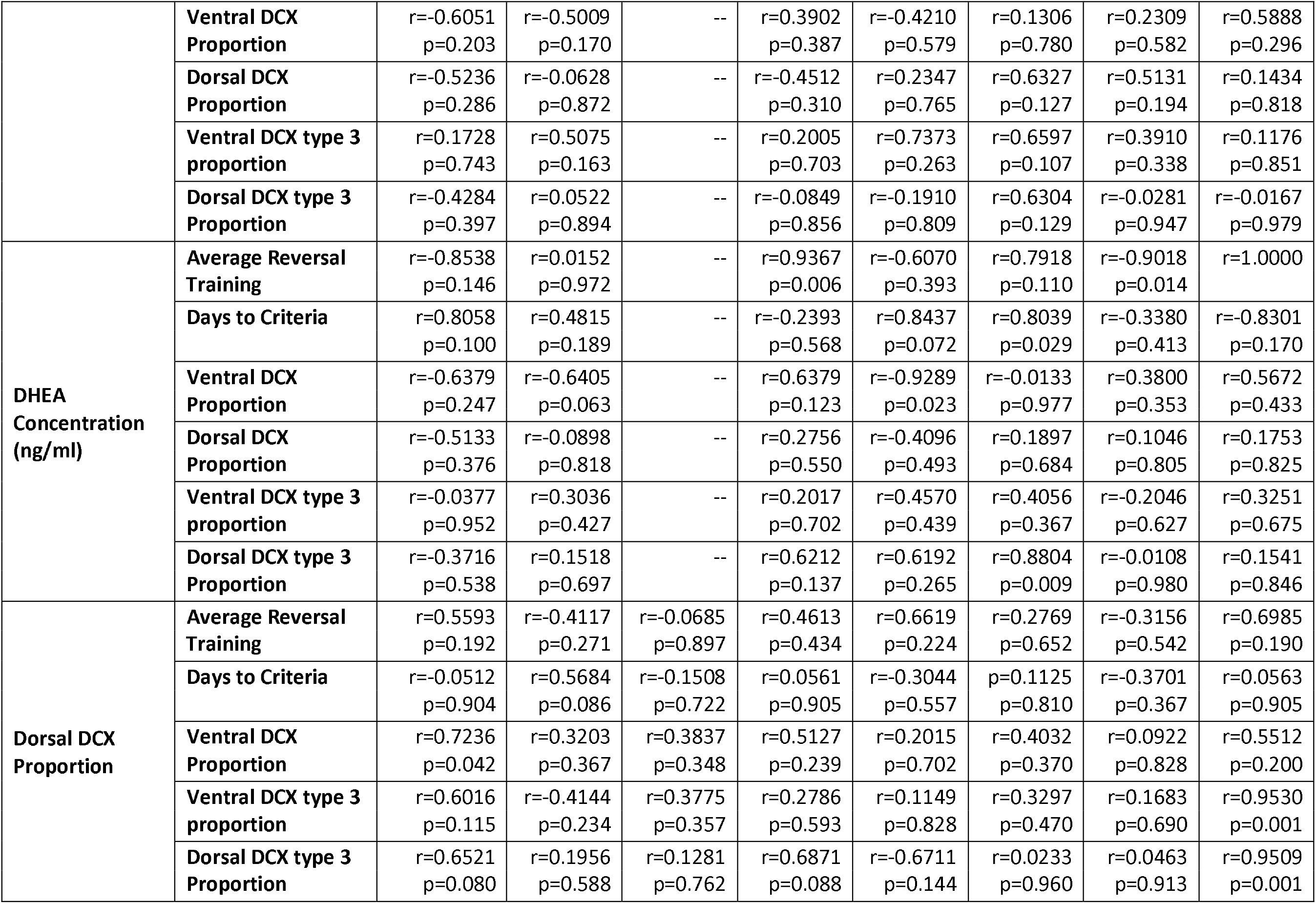
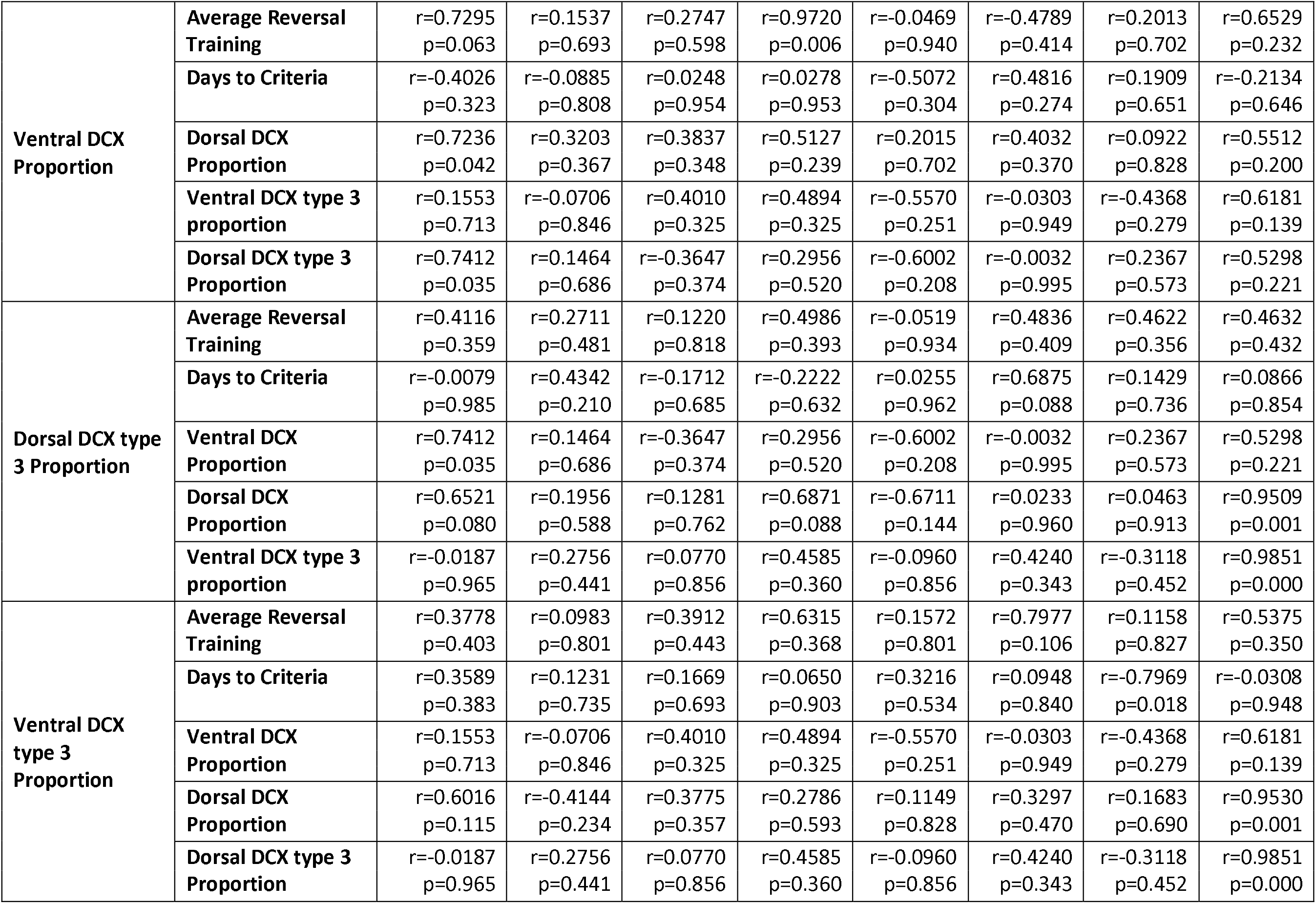
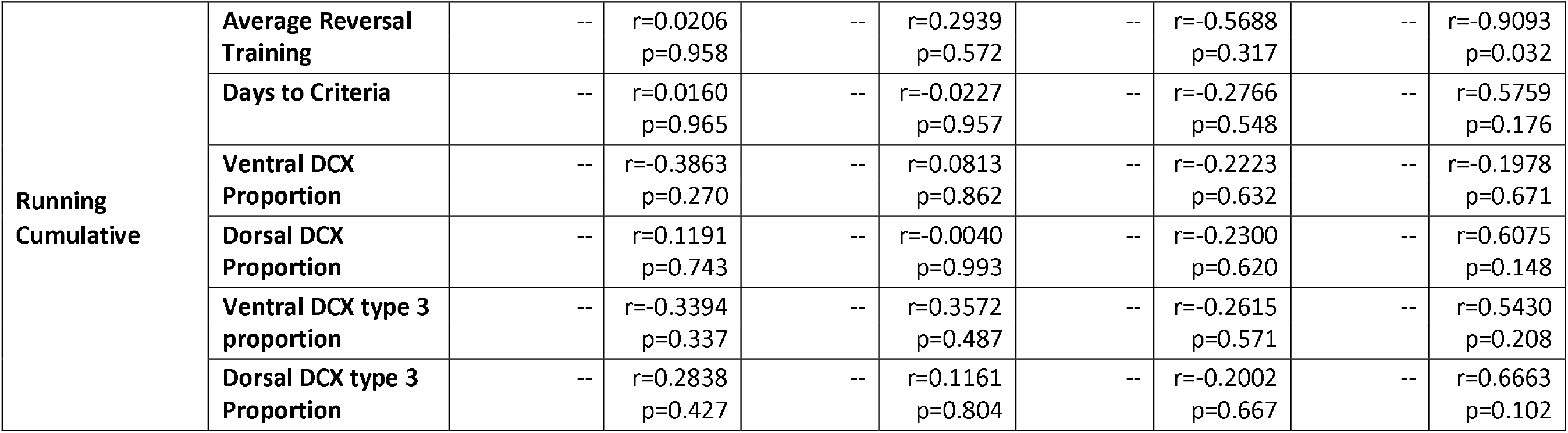
All Pearson product-moment correlations by genotype (Val/Val, Met/Met), sex (males, females), and exercise group (sedentary control (control), aerobic training (AT). DCX=Doublecortin expressing cells, DHEA= Dehydroepiandrosterone.

