## Supplemental Section for "Sex and BDNF Val66Met Polymorphism matter for exercise-induced increase in neurogenesis and cognition in middle-aged mice"

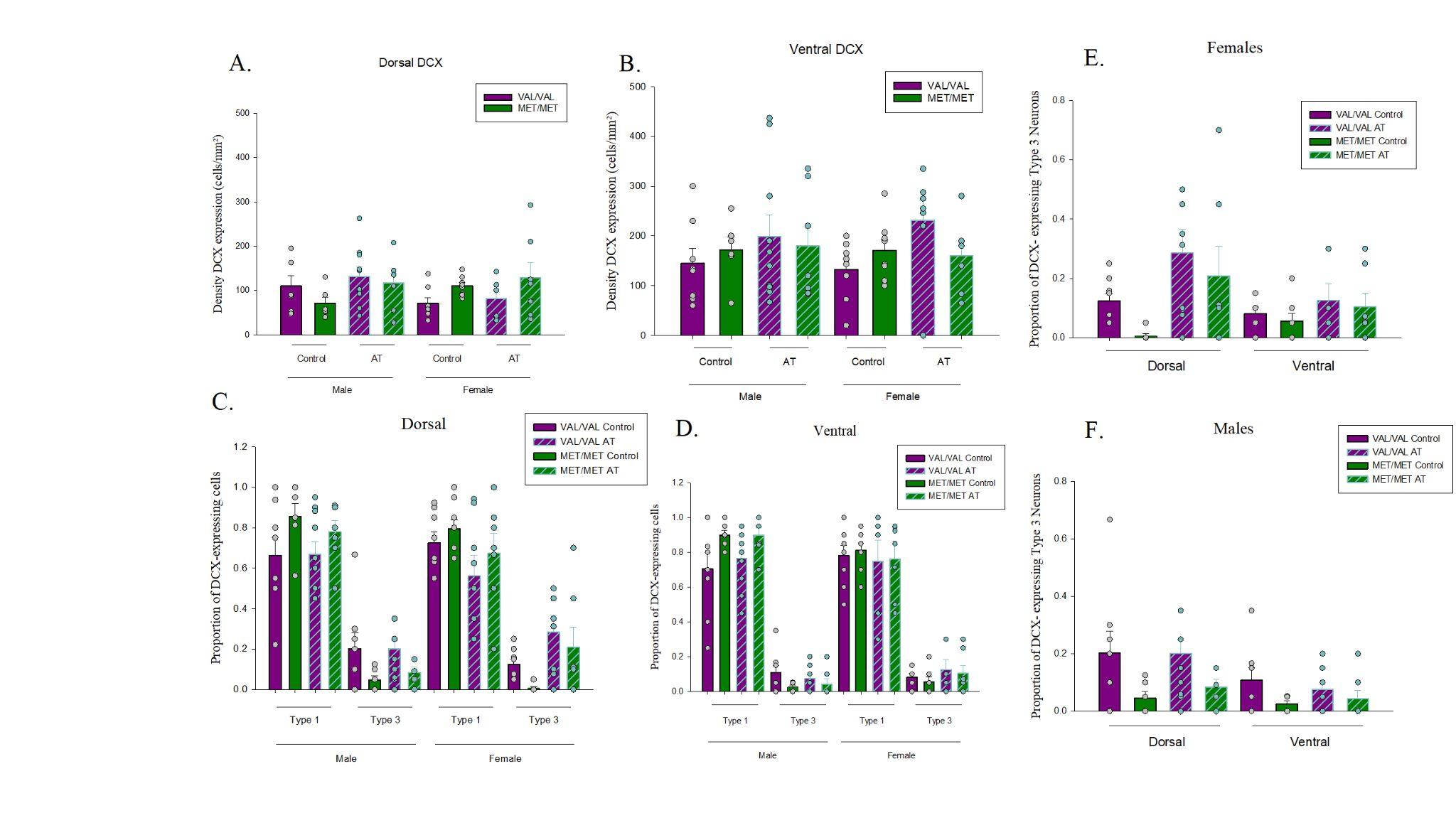

Supplemental Figure 1. Main findings separated by all groups. Dots represent individual data points. A-B) Density (cells/mm2) of doublecortin (DCX)-expressing cells by BDNF genotype, sex, and aerobic training (AT) in the dorsal (A) and ventral (B) region of the dentate gyrus. C-D) Maturity of DCX-expressing cells (type 1, and type 3) by BDNF genotype, sex, and AT in the dorsal (C) and the ventral (D) region of the dentate gyrus. E-F) The proportion of mature type 3 DCX-expressing cells by BDNF genotype, region, and AT in female (E) and male (F) mice.
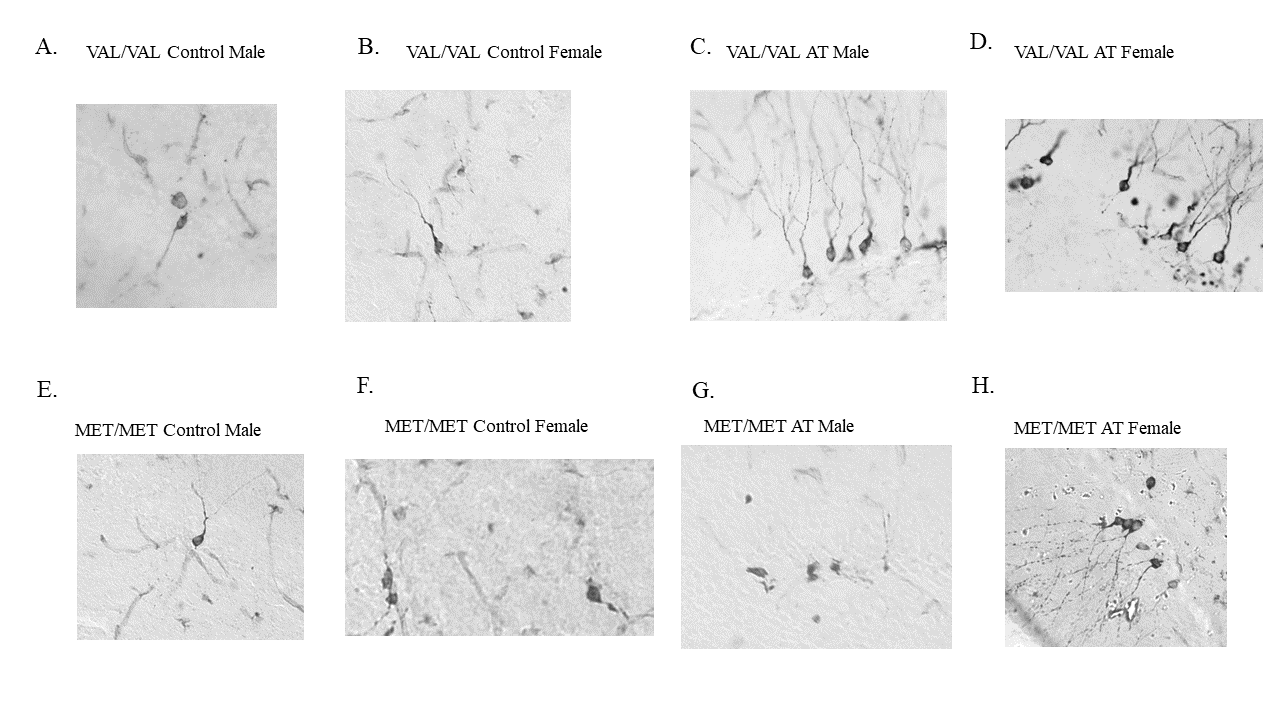

Supplemental Figure 2. Photomicrographs of doublecortin (DCX) expressing cells at 40x objective. A-D) BDNF VAL/VAL mice. E-H. BDNF MET/MET mice. A) Photomicrograph of a BDNF VAL/VAL control male mouse. B) Photomicrograph of a BDNF VAL/VAL control female mouse. C) Photomicrograph of a BDNF VAL/VAL aerobic training (AT) male mouse. D) Photomicrograph of a BDNF VAL/VAL AT female mouse. E) Photomicrograph of a BDNF MET/MET control male. F) Photomicrograph of a BDNF MET/MET control female. G) Photomicrograph of a BDNF MET/MET AT male. H) Photomicrograph of a BDNF MET/MET AT female.

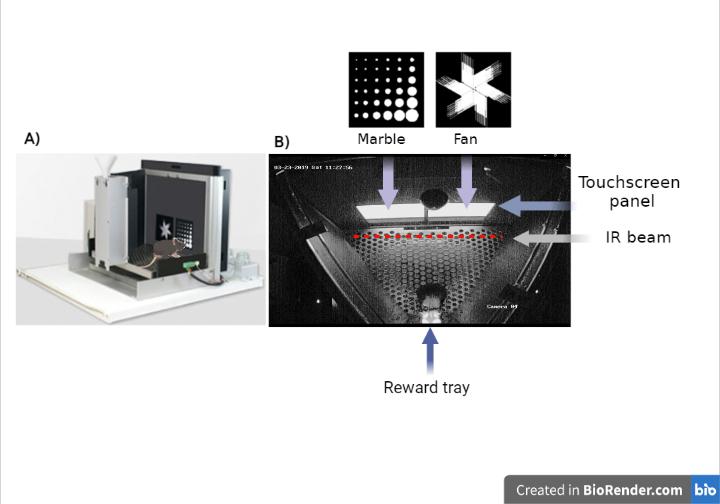

Supplemental Figure 3. A) Illustrated side view of the touchscreen chamber (modified from: edspace.american.edu/openbehavior/project/touchscreen-cognition-mousebytes/). Panel B) shows a real image from above of the trapezoidal chamber and the touchscreen panel that displays the stimulus (upper) and the reward tray (bottom).

Supplemental Table 1- All Pearson product-moment correlations by genotype (Val/Val, Met/Met), sex (males, females), and exercise group (sedentary control (control), aerobic training (AT). DCX = Doublecortin expressing cells, DHEA= Dehydroepiandrosterone.

|  | | **Val/Val** | | | | **Met/Met** | | | |
| --- | --- | --- | --- | --- | --- | --- | --- | --- | --- |
|  |  | **Male Control** | **Male AT** | **Female control** | **Female AT** | **Male Control** | **Male At** | **Female Control** | **Female AT** |
| **Adrenal Weight** | **Average Reversal** | r=0.0058 | r=-0.7809 | r=-0.4182 | r=-0.0342 | r=0.2501 | r=-0.1146 | r=-0.3255 | r=-0.1267 |
|  | **Training** | p=0.993 | p=0.067 | p=0.725 | p=0.966 | p=0.685 | p=0.854 | p=0.529 | p=0.839 |
|  | **Days to Criteria** | r=0.1377 | r=0.6521 | r=-0.6841 | r=-0.7130 | r=-0.0858 | r=-0.8670 | r=0.1309 | r=0.7251 |
|  |  | p=0.825 | p=0.160 | p=0.316 | p=0.112 | p=0.872 | p=0.012 | p=0.757 | p=0.065 |
|  | **Ventral DCX** | r=-0.3719 | r=-0.5523 | r=0.2413 | r=-0.1707 | r=0.3237 | r=-0.1760 | r=0.1067 | r=0.0375 |
|  | **Proportion** | p=0.538 | p=0.256 | p=0.759 | p=0.784 | p=0.531 | p=0.706 | p=0.801 | p=0.936 |
| **(g)** | **Dorsal DCX** | r=0.2249 | r=0.3844 | r=-0.5790 | r=-0.9038 | r=0.8659 | r=0.0336 | r=0.7105 | r=0.0905 |
|  | **Proportion** | p=0.716 | p=0.452 | p=0.421 | p=0.035 | p=0.026 | p=0.943 | p=0.048 | p=0.847 |
|  | **Ventral DCX type 3** | r=0.3324 | r=-0.3239 | r=-0.9705 | r=0.0605 | r=-0.1628 | r=0.1531 | r=-0.2932 | r=0.0333 |
|  | **proportion** | p=0.585 | p=0.531 | p=0.029 | p=0.940 | p=0.758 | p=0.743 | p=0.481 | p=0.944 |
|  | **Dorsal DCX type 3** | r=-0.2120 | r=-0.7240 | r=-0.1451 | r=-0.9156 | r=-0.7037 | r=-0.7168 | r=-0.1548 | r=0.1535 |
|  | **Proportion** | p=0.732 | p=0.104 | p=0.855 | p=0.029 | p=0.119 | p=0.070 | p=0.714 | p=0.742 |
| **Estradiol** | **Average Reversal** | r=0.0645 | r=0.1539 | -- | r=-0.3718 | r=0.1884 | r=0.3803 | r=-0.4283 | r=0.3081 |
|  | **Training** | p=0.935 | p=0.716 |  | p=0.468 | p=0.812 | p=0.528 | p=0.397 | p=0.801 |
|  | **Days to Criteria** | r=-0.4891 | r=-0.5751 | -- | r=0.0845 | r=0.4887 | r=0.7586 | r=-0.6387 | r=-0.3323 |
|  |  | p=0.403 | p=0.105 |  | p=0.842 | p=0.404 | p=0.048 | p=0.088 | p=0.585 |
|  | **Ventral DCX** | r=-0.1923 | r=0.5798 | -- | r=-0.3130 | r=-0.3535 | r=0.3241 | r=-0.0047 | r=0.1507 |
|  | **Proportion** | p=0.757 | p=0.102 |  | p=0.494 | p=0.559 | p=0.478 | p=0.991 | p=0.809 |
| **Concentration** |  |  |  |  |  |  |  |  |  |
|  | **Dorsal DCX** | r=-0.3201 | r=-0.3336 | -- | r=0.3046 | r=0.7198 | r=0.3968 | r=0.4025 | r=-0.1710 |
| **(ng/ml)** |  |  |  |  |  |  |  |  |  |
|  | **Proportion** | p=0.599 | p=0.380 |  | p=0.507 | p=0.170 | p=0.378 | p=0.323 | p=0.783 |
|  | **Ventral DCX type 3** | r=-0.4118 | r=0.2380 | -- | r=-0.3213 | r=0.5153 | r=-0.1262 | r=0.3854 | r=-0.3651 |
|  | **proportion** | p=0.491 | p=0.537 |  | p=0.535 | p=0.374 | p=0.787 | p=0.346 | p=0.546 |
|  | **Dorsal DCX type 3** | r=-0.0798 | r=-0.0846 | -- | r=0.0317 | r=-0.5241 | r=0.2708 | r=0.2116 | r=-0.3866 |
|  | **Proportion** | p=0.899 | p=0.829 |  | p=0.946 | p=0.365 | p=0.557 | p=0.615 | p=0.520 |
| **Progesterone Concentration** | **Average Reversal Training** | r=-0.3715 p=0.538 | r=0.1520 p=0.719 | -- | r=0.1129 p=0.831 | r=-0.1123 p=0.888 | r=0.5793 p=0.306 | r=-0.5872 p=0.220 | r=0.9555 p=0.191 |
|  | **Days to Criteria** | r=0.7984 | r=0.1373 | -- | r=0.5346 | r=0.3543 | r=0.4460 | r=-0.8271 | r=-0.7547 |
| **(ng/ml)** |  |  |  |  |  |  |  |  |  |
|  |  | p=0.057 | p=0.725 |  | p=0.172 | p=0.646 | p=0.316 | p=0.011 | p=0.140 |

|  | **Ventral DCX Proportion** | r=-0.6051 p=0.203 | r=-0.5009 p=0.170 | -- | r=0.3902 p=0.387 | r=-0.4210 p=0.579 | r=0.1306 p=0.780 | r=0.2309 p=0.582 | r=0.5888 p=0.296 |
| --- | --- | --- | --- | --- | --- | --- | --- | --- | --- |
|  | **Dorsal DCX Proportion** | r=-0.5236 p=0.286 | r=-0.0628 p=0.872 | -- | r=-0.4512 p=0.310 | r=0.2347 p=0.765 | r=0.6327 p=0.127 | r=0.5131 p=0.194 | r=0.1434 p=0.818 |
|  | **Ventral DCX type 3 proportion** | r=0.1728 p=0.743 | r=0.5075 p=0.163 | -- | r=0.2005 p=0.703 | r=0.7373 p=0.263 | r=0.6597 p=0.107 | r=0.3910 p=0.338 | r=0.1176 p=0.851 |
|  | **Dorsal DCX type 3 Proportion** | r=-0.4284 p=0.397 | r=0.0522 p=0.894 | -- | r=-0.0849 p=0.856 | r=-0.1910 p=0.809 | r=0.6304 p=0.129 | r=-0.0281 p=0.947 | r=-0.0167 p=0.979 |
| **DHEA** | **Average Reversal** | r=-0.8538 | r=0.0152 | -- | r=0.9367 | r=-0.6070 | r=0.7918 | r=-0.9018 | r=1.0000 |
|  | **Training** | p=0.146 | p=0.972 |  | p=0.006 | p=0.393 | p=0.110 | p=0.014 |  |
|  | **Days to Criteria** | r=0.8058 | r=0.4815 | -- | r=-0.2393 | r=0.8437 | r=0.8039 | r=-0.3380 | r=-0.8301 |
|  |  | p=0.100 | p=0.189 |  | p=0.568 | p=0.072 | p=0.029 | p=0.413 | p=0.170 |
|  | **Ventral DCX** | r=-0.6379 | r=-0.6405 | -- | r=0.6379 | r=-0.9289 | r=-0.0133 | r=0.3800 | r=0.5672 |
|  | **Proportion** | p=0.247 | p=0.063 |  | p=0.123 | p=0.023 | p=0.977 | p=0.353 | p=0.433 |
| **Concentration** |  |  |  |  |  |  |  |  |  |
|  | **Dorsal DCX** | r=-0.5133 | r=-0.0898 | -- | r=0.2756 | r=-0.4096 | r=0.1897 | r=0.1046 | r=0.1753 |
| **(ng/ml)** |  |  |  |  |  |  |  |  |  |
|  | **Proportion** | p=0.376 | p=0.818 |  | p=0.550 | p=0.493 | p=0.684 | p=0.805 | p=0.825 |
|  | **Ventral DCX type 3** | r=-0.0377 | r=0.3036 | -- | r=0.2017 | r=0.4570 | r=0.4056 | r=-0.2046 | r=0.3251 |
|  | **proportion** | p=0.952 | p=0.427 |  | p=0.702 | p=0.439 | p=0.367 | p=0.627 | p=0.675 |
|  | **Dorsal DCX type 3** | r=-0.3716 | r=0.1518 | -- | r=0.6212 | r=0.6192 | r=0.8804 | r=-0.0108 | r=0.1541 |
|  | **Proportion** | p=0.538 | p=0.697 |  | p=0.137 | p=0.265 | p=0.009 | p=0.980 | p=0.846 |
| **Dorsal DCX** | **Average Reversal** | r=0.5593 | r=-0.4117 | r=-0.0685 | r=0.4613 | r=0.6619 | r=0.2769 | r=-0.3156 | r=0.6985 |
|  | **Training** | p=0.192 | p=0.271 | p=0.897 | p=0.434 | p=0.224 | p=0.652 | p=0.542 | p=0.190 |
|  | **Days to Criteria** | r=-0.0512 | r=0.5684 | r=-0.1508 | r=0.0561 | r=-0.3044 | p=0.1125 | r=-0.3701 | r=0.0563 |
|  |  | p=0.904 | p=0.086 | p=0.722 | p=0.905 | p=0.557 | p=0.810 | p=0.367 | p=0.905 |
|  | **Ventral DCX** | r=0.7236 | r=0.3203 | r=0.3837 | r=0.5127 | r=0.2015 | r=0.4032 | r=0.0922 | r=0.5512 |
| **Proportion** | **Proportion** | p=0.042 | p=0.367 | p=0.348 | p=0.239 | p=0.702 | p=0.370 | p=0.828 | p=0.200 |
|  | **Ventral DCX type 3** | r=0.6016 | r=-0.4144 | r=0.3775 | r=0.2786 | r=0.1149 | r=0.3297 | r=0.1683 | r=0.9530 |
|  | **proportion** | p=0.115 | p=0.234 | p=0.357 | p=0.593 | p=0.828 | p=0.470 | p=0.690 | p=0.001 |
|  | **Dorsal DCX type 3** | r=0.6521 | r=0.1956 | r=0.1281 | r=0.6871 | r=-0.6711 | r=0.0233 | r=0.0463 | r=0.9509 |
|  | **Proportion** | p=0.080 | p=0.588 | p=0.762 | p=0.088 | p=0.144 | p=0.960 | p=0.913 | p=0.001 |

| **Ventral DCX** | **Average Reversal** | r=0.7295 | r=0.1537 | r=0.2747 | r=0.9720 | r=-0.0469 | r=-0.4789 | r=0.2013 | r=0.6529 |
| --- | --- | --- | --- | --- | --- | --- | --- | --- | --- |
|  | **Training** | p=0.063 | p=0.693 | p=0.598 | p=0.006 | p=0.940 | p=0.414 | p=0.702 | p=0.232 |
|  | **Days to Criteria** | r=-0.4026 | r=-0.0885 | r=0.0248 | r=0.0278 | r=-0.5072 | r=0.4816 | r=0.1909 | r=-0.2134 |
|  |  | p=0.323 | p=0.808 | p=0.954 | p=0.953 | p=0.304 | p=0.274 | p=0.651 | p=0.646 |
|  | **Dorsal DCX** | r=0.7236 | r=0.3203 | r=0.3837 | r=0.5127 | r=0.2015 | r=0.4032 | r=0.0922 | r=0.5512 |
| **Proportion** | **Proportion** | p=0.042 | p=0.367 | p=0.348 | p=0.239 | p=0.702 | p=0.370 | p=0.828 | p=0.200 |
|  | **Ventral DCX type 3** | r=0.1553 | r=-0.0706 | r=0.4010 | r=0.4894 | r=-0.5570 | r=-0.0303 | r=-0.4368 | r=0.6181 |
|  | **proportion** | p=0.713 | p=0.846 | p=0.325 | p=0.325 | p=0.251 | p=0.949 | p=0.279 | p=0.139 |
|  | **Dorsal DCX type 3** | r=0.7412 | r=0.1464 | r=-0.3647 | r=0.2956 | r=-0.6002 | r=-0.0032 | r=0.2367 | r=0.5298 |
|  | **Proportion** | p=0.035 | p=0.686 | p=0.374 | p=0.520 | p=0.208 | p=0.995 | p=0.573 | p=0.221 |
| **Dorsal DCX type** | **Average Reversal** | r=0.4116 | r=0.2711 | r=0.1220 | r=0.4986 | r=-0.0519 | r=0.4836 | r=0.4622 | r=0.4632 |
|  | **Training** | p=0.359 | p=0.481 | p=0.818 | p=0.393 | p=0.934 | p=0.409 | p=0.356 | p=0.432 |
|  | **Days to Criteria** | r=-0.0079 | r=0.4342 | r=-0.1712 | r=-0.2222 | r=0.0255 | r=0.6875 | r=0.1429 | r=0.0866 |
|  |  | p=0.985 | p=0.210 | p=0.685 | p=0.632 | p=0.962 | p=0.088 | p=0.736 | p=0.854 |
|  | **Ventral DCX** | r=0.7412 | r=0.1464 | r=-0.3647 | r=0.2956 | r=-0.6002 | r=-0.0032 | r=0.2367 | r=0.5298 |
| **3 Proportion** | **Proportion** | p=0.035 | p=0.686 | p=0.374 | p=0.520 | p=0.208 | p=0.995 | p=0.573 | p=0.221 |
|  | **Dorsal DCX** | r=0.6521 | r=0.1956 | r=0.1281 | r=0.6871 | r=-0.6711 | r=0.0233 | r=0.0463 | r=0.9509 |
|  | **Proportion** | p=0.080 | p=0.588 | p=0.762 | p=0.088 | p=0.144 | p=0.960 | p=0.913 | p=0.001 |
|  | **Ventral DCX type 3** | r=-0.0187 | r=0.2756 | r=0.0770 | r=0.4585 | r=-0.0960 | r=0.4240 | r=-0.3118 | r=0.9851 |
|  | **proportion** | p=0.965 | p=0.441 | p=0.856 | p=0.360 | p=0.856 | p=0.343 | p=0.452 | p=0.000 |
| **Ventral DCX** | **Average Reversal** | r=0.3778 | r=0.0983 | r=0.3912 | r=0.6315 | r=0.1572 | r=0.7977 | r=0.1158 | r=0.5375 |
|  | **Training** | p=0.403 | p=0.801 | p=0.443 | p=0.368 | p=0.801 | p=0.106 | p=0.827 | p=0.350 |
|  | **Days to Criteria** | r=0.3589 | r=0.1231 | r=0.1669 | r=0.0650 | r=0.3216 | r=0.0948 | r=-0.7969 | r=-0.0308 |
|  |  | p=0.383 | p=0.735 | p=0.693 | p=0.903 | p=0.534 | p=0.840 | p=0.018 | p=0.948 |
|  | **Ventral DCX** | r=0.1553 | r=-0.0706 | r=0.4010 | r=0.4894 | r=-0.5570 | r=-0.0303 | r=-0.4368 | r=0.6181 |
| **type 3** |  |  |  |  |  |  |  |  |  |
|  | **Proportion** | p=0.713 | p=0.846 | p=0.325 | p=0.325 | p=0.251 | p=0.949 | p=0.279 | p=0.139 |
| **Proportion** |  |  |  |  |  |  |  |  |  |
|  | **Dorsal DCX** | r=0.6016 | r=-0.4144 | r=0.3775 | r=0.2786 | r=0.1149 | r=0.3297 | r=0.1683 | r=0.9530 |
|  | **Proportion** | p=0.115 | p=0.234 | p=0.357 | p=0.593 | p=0.828 | p=0.470 | p=0.690 | p=0.001 |
|  | **Dorsal DCX type 3** | r=-0.0187 | r=0.2756 | r=0.0770 | r=0.4585 | r=-0.0960 | r=0.4240 | r=-0.3118 | r=0.9851 |
|  | **Proportion** | p=0.965 | p=0.441 | p=0.856 | p=0.360 | p=0.856 | p=0.343 | p=0.452 | p=0.000 |

| **Running** | **Average Reversal** | -- | r=0.0206 | -- | r=0.2939 | -- | r=-0.5688 | -- | r=-0.9093 |
| --- | --- | --- | --- | --- | --- | --- | --- | --- | --- |
|  | **Training** |  | p=0.958 |  | p=0.572 |  | p=0.317 |  | p=0.032 |
|  | **Days to Criteria** | -- | r=0.0160 | -- | r=-0.0227 | -- | r=-0.2766 | -- | r=0.5759 |
|  |  |  | p=0.965 |  | p=0.957 |  | p=0.548 |  | p=0.176 |
|  | **Ventral DCX** | -- | r=-0.3863 | -- | r=0.0813 | -- | r=-0.2223 | -- | r=-0.1978 |
|  | **Proportion** |  | p=0.270 |  | p=0.862 |  | p=0.632 |  | p=0.671 |
| **Cumulative** | **Dorsal DCX** | -- | r=0.1191 | -- | r=-0.0040 | -- | r=-0.2300 | -- | r=0.6075 |
|  | **Proportion** |  | p=0.743 |  | p=0.993 |  | p=0.620 |  | p=0.148 |
|  | **Ventral DCX type 3** | -- | r=-0.3394 | -- | r=0.3572 | -- | r=-0.2615 | -- | r=0.5430 |
|  | **proportion** |  | p=0.337 |  | p=0.487 |  | p=0.571 |  | p=0.208 |
|  | **Dorsal DCX type 3** | -- | r=0.2838 | -- | r=0.1161 | -- | r=-0.2002 | -- | r=0.6663 |
|  | **Proportion** |  | p=0.427 |  | p=0.804 |  | p=0.667 |  | p=0.102 |
